# Annotation of the non-canonical translatome reveals that CHO cell microproteins are a new class of mAb drug product impurity

**DOI:** 10.1101/2022.01.20.475618

**Authors:** Marina Castro-Rivadeneyra, Ioanna Tzani, Paul Kelly, Lisa Strasser, Felipe Guapo, Ciara Tierney, Michelle Chain, Lin Zhang, Martin Clynes, Barry L. Karger, Niall Barron, Jonathan Bones, Colin Clarke

## Abstract

Chinese hamster ovary (CHO) cells are used to produce almost 90% of therapeutic monoclonal antibodies (mAbs). The annotation of non-canonical translation events in these cellular factories remains incomplete, limiting not only our ability to study CHO cell biology but also detect host cell protein (HCP) contaminants in the final mAb drug product. We utilised ribosome footprint profiling (Ribo-seq) to identify novel open reading frames (ORFs) including N-terminal extensions and thousands of short ORFs (sORFs) predicted to encode microproteins. Mass spectrometry-based HCP analysis of four commercial mAb drug products using the extended protein sequence database revealed the presence of microprotein impurities for the first time. We also show that microprotein abundance varies with growth phase and can be affected by the cell culture environment. In addition, our work provides a vital resource to facilitate future studies of non-canonical translation as well as the regulation of protein synthesis in CHO cell lines.

## 1. Introduction

Chinese hamster ovary (CHO) cells are the predominant mammalian expression host for the production of biologics, with nearly 90% of therapeutic monoclonal antibodies (mAbs) produced in this cell line^1^. During the “upstream” cell culture phase of production, CHO cells continually secrete a recombinant mAb into the supernatant. Upon completion, a series of “downstream” purification steps are required to recover the product in the harvested cell culture fluid and reduce a range of impurities including those originating from the host CHO cell line. Host cell proteins (HCPs) present in the final drug product are a particular concern, due to the risk that a HCP could elicit an immune response in the patient or reduce efficacy^2^. In addition, the presence of proteolytic HCPs can degrade or affect the stability of the mAb^3,4^. Regulatory authorities consider the amount of HCP in the final product to be a critical quality attribute, and require that the total HCP concentration be < 100 ppm^5^.

The efficacy and safety track record of therapeutic proteins produced in CHO cells is testament to the continual commitment of biopharmaceutical companies and regulatory authorities to ensuring product quality. Enzyme-linked immunosorbent assays (ELISA) that enable sensitive total HCP quantitation are widely used in batch release during commercial manufacturing^6^. It is recognised, however, that this method can be limited in terms of coverage^7^. For instance, some low molecular weight proteins^7^ may be weakly immunogenic or indeed a particular protein may not elicit a response in the species immunised to generate the HCP assay^8^. Regulatory authorities now recommend^7^ the use of mass spectrometry (MS) as an orthogonal HCP detection method^5^ to enable the identification and quantification of individual HCPs, even those at low concentrations. The resulting data can be used during process development, to monitor the HCP contaminants present at each unit operation of a downstream purification process as well as demonstrate HCP clearance from the final drug product^7,9^. Identification of HCPs can also be used to guide upstream process development^10^, or identify targets for cell line engineering to remove unwanted HCPs^11^.

Since publication of the first CHO cell genome^12^ and of CHO cell-specific protein databases^13^ the detection of CHO HCP impurities in mAb drug product using MS has significantly improved. The quality of available genomes has steadily improved over time, and, with the release of the Chinese hamster PICRH genome, the field now has a reference assembly comparable to that of model organisms^14^. While annotation of the transcriptome has progressed significantly, characterisation of the proteome is more challenging and remains incomplete, therefore limiting the ability of MS to detect the complete spectrum of potential HCP impurities.

The Chinese hamster reference genome has predominantly been annotated *via* a combination of *ab initio* computational pipelines, homology, ESTs, and transcriptomics data. The Lewis lab elegantly demonstrated the effectiveness of ribosome footprint profiling (Ribo-seq) to identify translated regions of the Chinese hamster genome^15^. Ribo-seq enables transcriptome-wide determination of ribosome occupancy at single nucleotide resolution enabling open reading frame (ORF) annotation, and, when combined with RNA-seq, variations in translational regulation^16^. The technique utilises chemical or physical inhibitors to arrest translation and fixes translating ribosomes in position resulting in the protection of ∼30 nt of mRNA within the ribosome from subsequent enzymatic degradation. The resulting monosomes are purified via sucrose gradient, sucrose cushion or size exclusion chromatography, followed by the isolation of ribosome protected fragments (RPFs) through size selection, from which a sequencing library is prepared. Sequencing of RPFs and alignment to a reference genome or transcriptome permits the identification and quantitation of regions undergoing active translation. Over the last decade Ribo-seq has provided compelling evidence that the traditional rules of eukaryotic translation need to be revised. For example, translation initiation at near-cognate codons (CUG, GUG, UUG) is more widespread in mammalian genomes than previously thought^17^.

Data from Ribo-seq experiments has been used to annotate a range of non-canonical ORFs^18^, including N-terminal extensions^19^, detect translation of RNAs previously thought to be non-coding^20^, and study the regulatory role of ORFs in the 5’ leader sequence of mRNAs (i.e., upstream open reading frames)^21^. Ribo-seq has also revealed the existence of small open reading frames (sORFs) that produce potentially functional microproteins (classified as proteins < 100 aa) in a diverse range of organisms including *Drosophila*^22^, zebrafish^23^, mouse^24^ and human^25,26^. Studies have so far shown that microproteins are involved in a variety of cellular processes such as oxidative phosphorylation^27^, mitochondrial translation^28^, metabolism^29^, DNA repair^30^ and can also act as transcription factors^31^.

There has been considerable interest in enhancing the efficiency of CHO cell factories for mAb production using systems biology^32^ and cell line engineering^33^. Yet, while the importance of non-canonical ORFs is becoming increasingly understood in other organisms, the study of their role in CHO cell biology is severely restricted by the lack of annotation. Perhaps more surprisingly, given the fundamental role of protein synthesis in mAb production, there is a lack of knowledge of how translational regulation impacts CHO cell behaviour during cell culture processes for mAb manufacturing. Indeed, apart from a small number of studies^15,34^, Ribo-seq has not received widespread attention in the field and the capability of the technique to study CHO cell translation regulation has yet to be demonstrated. An improvement in the annotation of non-canonical ORFs in Chinese hamster genome and transcriptome-wide analysis of protein synthesis in CHO cells has the potential to identify new avenues to improve recombinant therapeutic protein production processes.

In this study, we conducted a series of Ribo-seq experiments employing different translation inhibitors to analyse translation elongation, and crucially translation initiation for the first time in CHO cells. Using these data, we have significantly enhanced the annotation of non-canonical ORFs in the Chinese hamster genome. We identified novel translation events in previously annotated protein-coding genes as well as thousands of new short ORFs predicted to encode microproteins. Our work has improved MS-based HCP detection in mAb drug products and provides an essential foundation for future studies of non-canonical translation events and the control of protein synthesis in this important cell line.

## 2. Results

### 2.1 Transcriptome-wide analysis of CHO cell translation initiation and elongation using Ribo-seq

Ribo-seq and RNA-seq profiling was carried out on a small-scale cell culture model that simulates a “temperature shift” i.e., a transition to mildly hypothermic conditions once cells have reached sufficient density in the bioreactor that enhances the efficiency of some in industrial manufacturing processes ^35^. Previous studies in our laboratory using a mAb producing CHO K1 cell line (CHO K1-mAb) demonstrate that a reduction in cell culture temperature reduces the growth rate, changes the metabolism and results in significant differences in gene expression^36^. In this study, we reasoned that our temperature shift model would also induce widespread changes in translation and would provide an ideal opportunity to identify new CHO cell ORFs.

To capture the Ribo-seq data we conducted two identical cell culture experiments for the analysis of translation initiation and elongation. For both experiments, 8 replicate shake flasks were first grown for 48 hr at 37°C before the cell culture temperature was reduced to 31°C (temperature shifted (TS) group; n = 4) while maintaining the remainder at 37°C (non-temperature shift (NTS) group; n = 4). Samples from both the TS and NTS groups were harvested for Ribo-seq 24 hr after the reduction of cell culture temperature (Figure 1a) at which point there was an average of 30% lower (initiation experiment) and 24% (elongation experiment) in the cell density of the TS sample group (Supplementary Figure 1; Supplementary Data 1).

**Figure 1:**
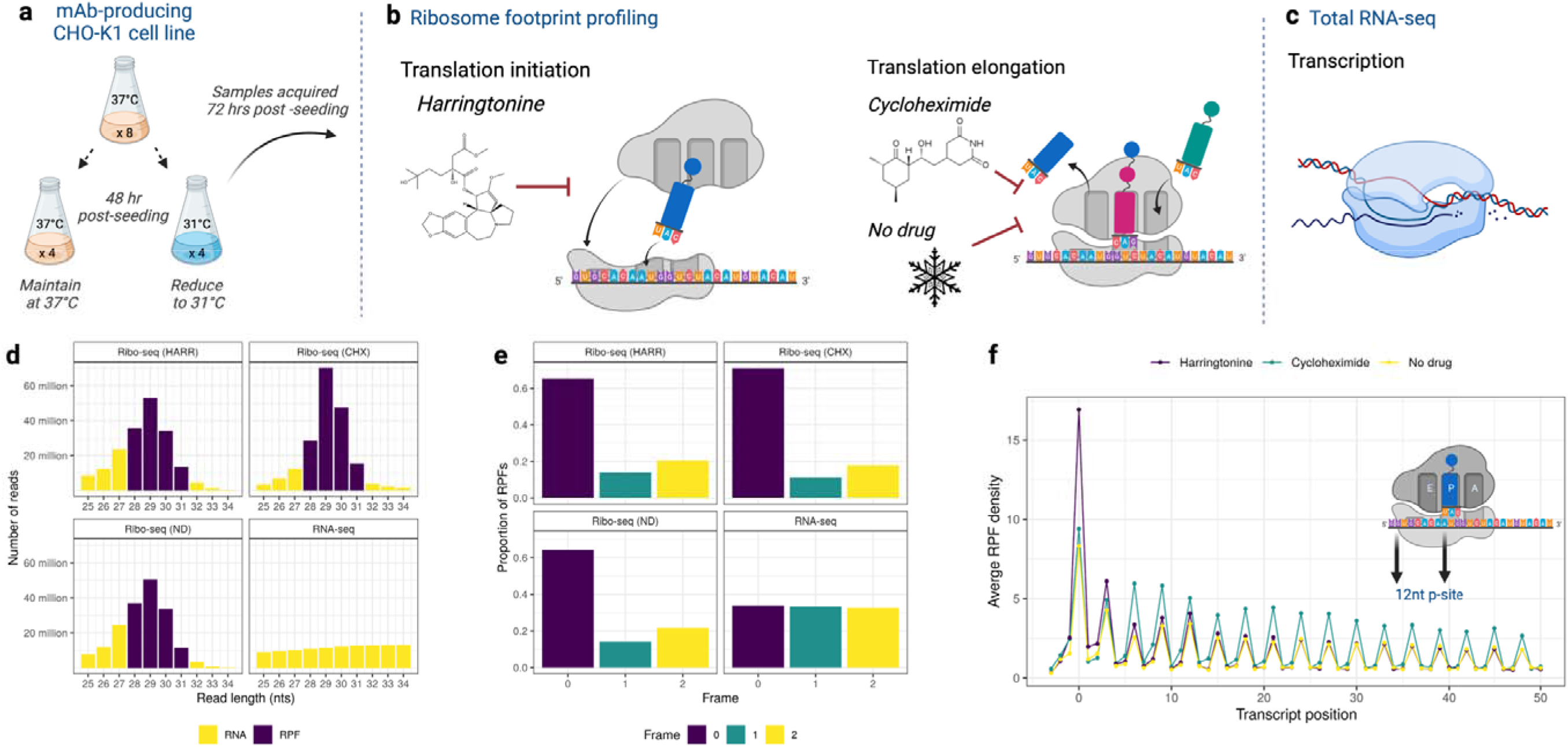
Analysis of CHO cell translation using ribosome footprint profiling. **(a)** 8 replicate shake flasks were seeded with a mAb producing CHO K1 cell line cultured for 48 hr, at this point the temperature of 4 shake flasks was reduced to 31°C. At 72 hr post-seeding, samples were harvested from the non-temperature and temperature shifted cultures. We utilised **(b)** Ribo-seq using different inhibitors to capture information from initiating (harringtonine) and elongating (cycloheximide and no drug) ribosomes. In addition, **(c)** RNA-seq was used to characterise the transcriptome. Following pre-processing of the raw Ribo-seq data, we **(d)** retained reads within the expected size range of RPFs. An optimum P-site offset of 12 nucleotides was selected for all datasets, where **(e)** an average of 60% of RPFs was found to exhibit the expected triplet periodicity. A metagene analysis was conducted for each Ribo-seq dataset, confirming **(f)** the expected enrichment of RPFs at the TIS of annotated protein coding genes in the harringtonine data when compared to the cycloheximide and no-drug treated data.

To capture a snapshot of the CHO cell translatome, we performed ribosome footprint profiling experiments using harringtonine (HARR) (n =8), an inhibitor of translation initiation^24^, and cycloheximide (CHX) (n = 8) an inhibitor of translation elongation (Figure 1b)^16^. For each harringtonine sample, a matched Ribo-seq sample (n = 8) was treated with DMSO and flash frozen to arrest translation (we refer to these data as “No-drug” (ND)). For the CHX samples, matched gene expression profiles were acquired using total RNA-seq (n = 8) (Figure 1c) to enable the identification of significant differences in translational efficiency (TE) between the NTS and TS sample groups.

Sequencing of the 24 resulting Ribo-seq libraries yielded an average of ∼68, ∼67 and ∼58 million per sample for the CHX, HARR and ND Ribo-seq, respectively while an average of ∼56 million reads per sample were obtained for the 8 RNA-seq libraries. Low quality reads were removed, and adapter sequences trimmed from the raw Ribo-seq and RNA-seq data (Supplementary Data 2). For Ribo-seq data, an additional filtering stage was carried out to eliminate contamination from non-coding RNA. Reads were mapped to STAR^37^ indices constructed from *Cricetulus griseus* rRNA, tRNA and snoRNA sequences obtained from v18 of the RNA Central database (The RNAcentral Consortium, 2019). Reads aligning to any of these indices were discarded from further analysis. This filtering stage removed an average of ∼40%, ∼46% and ∼50% of trimmed reads for the CHX, HARR and ND samples, respectively (Supplementary Figure 2; Supplementary Data 2).

Next, we examined the remaining Ribo-seq reads within the expected RPF length range (25-34nt) to select the P-site offset (the distance from the 5’ end of a read to the first nucleotide of the P-site codon) (Figure 1d). Each Ribo-seq dataset was mapped to the Chinese hamster PICRH-1.0 genome using STAR^37^. The Plastid tool^39^ was used to assess the P-site offset and subsequently determine the proportion of reads exhibiting triplet periodicity for NCBI-annotated canonical protein coding genes for each offset. Following this analysis, we retained the reads between 28-31 nt for further analysis (Figure 1e). The optimum P-site offset was found to be 12 nt, for which 60% of reads exhibited the expected triplet periodicity for each Ribo-seq dataset. Prior to *de novo* ORF identification, we confirmed the expected preferential enrichment of ribosomes at the translation initiation sites (TIS) of annotated protein coding genes for the HARR Ribo-seq data in comparison to the CHX and ND Ribo-seq data (Figure 1f).

### 2.2 Ribo-seq enables the characterisation of novel ORFs in the Chinese hamster genome

The Ribo-seq data was used to refine the annotation of translated regions of the Chinese hamster PICRH-1.0 genome by conducting a transcriptome-wide analysis using ORF-RATER^40^. The ORF-RATER algorithm integrates initiation and elongation Ribo-seq data to enable the identification of unannotated ORFs by first finding all potential ORFs beginning at user defined start codons that have an in-frame stop codon. The experimental Ribo-seq data is then used to confirm occupancy at each TIS and assess whether the putative ORF is undergoing active translation. To maximise the sensitivity of ORF detection, we merged the RPFs for all replicates in each type of Ribo-seq experiment yielding a total of approximately 136, 161 and 132 million RPFs for the harringtonine, cycloheximide, and no-drug treated Ribo-seq, respectively. Prior to ORF identification, we followed the ORF-RATER method and removed transcripts originating from pseudogenes (n = 4,583), transcripts with low RPF coverage (n = 19,357) and transcripts where the RPFs were mapped to a small number of positions (n = 1,538). For the remaining transcripts, the initial ORF-RATER search step was limited to ORFs that began at an AUG or near-cognate start codons (CUG, GUG and UUG). To determine if a potential TIS was occupied, only the RPF data from the HARR Ribo-seq was considered while CHX and ND-treated Ribo-seq data was utilised to determine if putative ORFs were translated by comparing the RPF occupancy of each ORF to the typical pattern of translation elongation observed for annotated Chinese hamster CDSs.

An initial group of 26,606 ORFs identified by ORF-RATER with an ORF-RATER score of ≥ 0.5^41,42^ and ORF length ≥ 5 aa was selected for further analysis. The proteoforms identified included those present in the current annotation of the Chinese hamster genome (i.e., Annotated) and N-terminal extensions (i.e., Extension). Two distinct classes of ORFs initiating upstream of the annotated CDS (i.e., the main ORF) were also identified. The first type, called upstream ORFs (i.e., uORFs) initiate upstream and terminate before the start codon of the main ORF. The second upstream ORF type, termed overlapping upstream open reading frames (ouORFs), also initiates in the 5’ leader of mRNAs but extends downstream beyond the start codon of the main ORF and is therefore translated in a different reading frame. We also identified ORFs in transcripts that had both unannotated start and stop codons in the PICRH-1.0 genome (“New” ORFs).

The conditions used to inhibit translation initiation can, in some cases, lead to the identification of false positive internal ORFs due to capture of residual elongating ribosomes^41^. In our case, we utilised flash freezing in combination with harringtonine, which will also result in the capture of a proportion of RPFs from elongating ribosomes, however, this will almost certainly lead to erroneous identifications. To reduce false positives from internal TIS, we excluded truncated ORF (n = 8,856) and internal ORF (n = 1,469) classifications from further analysis. In addition, where more than one upstream ORF (uORF), start overlapping uORF (ouORF) or “New” ORF had the same stop codon, we retained only the longest of these ORFs, resulting in the elimination of a further 941 ORFs. Following this stringent filtering process, 15,340 high confidence ORFs were retained (Figure 2a, Supplementary Data 3), with 49.3% (n = 7,769) of the identified ORFs not annotated in the Chinese hamster PICRH-1.0 genome. 58% of these new identifications start at near-cognate codons (i.e., CUG, GUG or UUG). The ability to identify initiation at non-AUG codons enabled us to identify alternative ORFs of conventional protein coding genes that would not be possible with previous annotation approaches for the Chinese hamster genome. For instance, 15.1% (n = 1,176) of novel ORFs were isoforms of annotated proteins and 12.1% (n = 941) were N-terminal extensions of annotated protein coding transcripts (e.g., Aurora kinase A (Figure 2b)).

**Figure 2:**
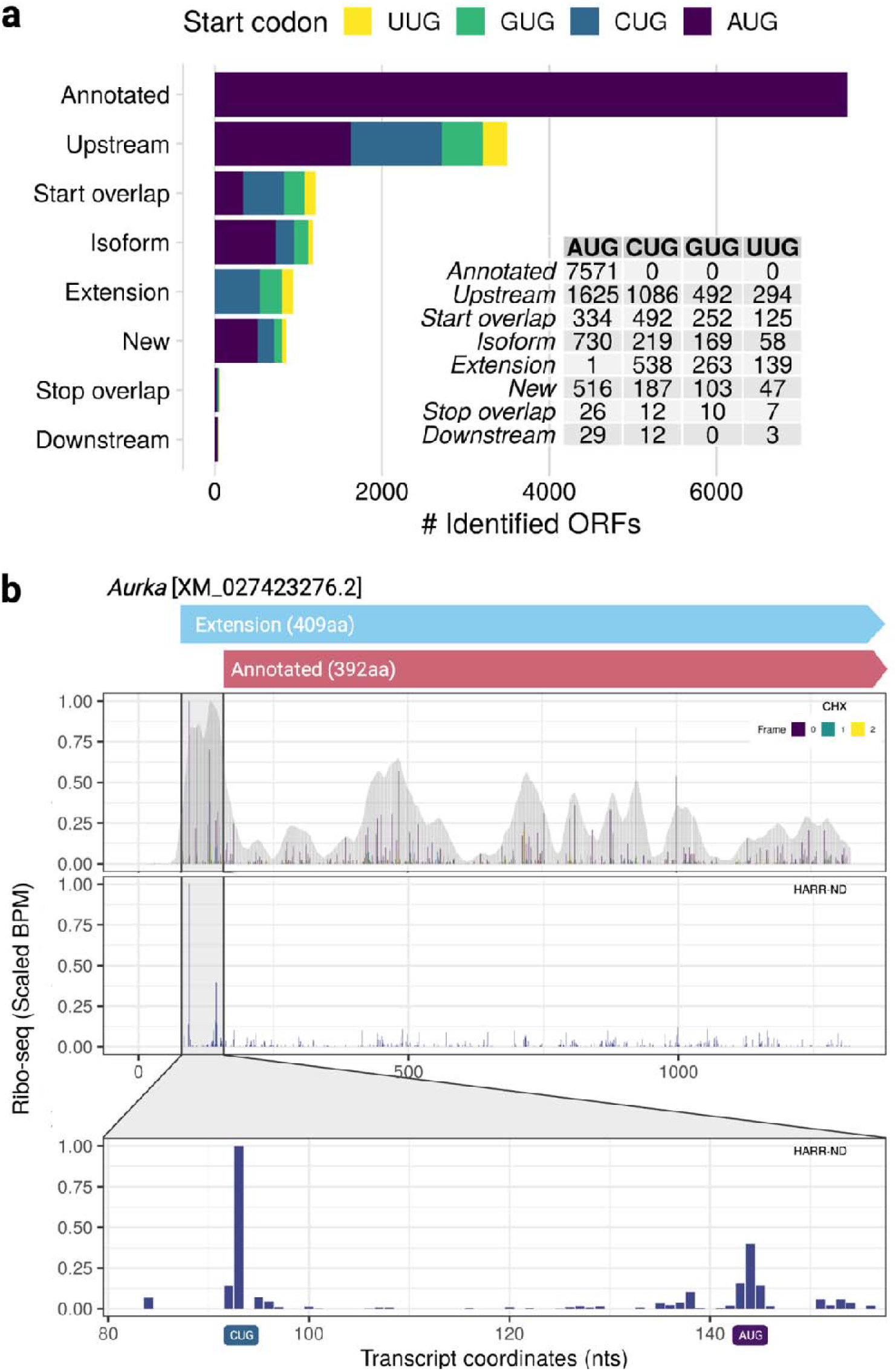
Ribo-seq identifies thousands of novel CHO cell ORFs. In this study, we utilised the ORF-RATER algorithm to identify ORFs initiated at near cognate (i.e., NUG) start codons from the Ribo-seq data. A total of **(a)** 15,340 ORFs were identified including 7,769 that were not previously annotated in the Chinese hamster genome. These new ORFs included N-terminal extensions of annotated protein coding genes. For instance, we identified a CUG initiated extension of **(b)** a transcript of the *Aurka* kinase gene. The CHX coverage of the transcript is shown (full coverage and P-site offset [coloured by reading frame relative to the annotated TIS]) along with the HARR-ND coverage (P-site offset) illustrating the initiation signal at the CUG start codon upstream of the NCBI annotated AUG start codon.

### 2.3 The Chinese hamster genome harbours thousands of short open reading frames

The ORF-RATER algorithm also identified thousands of previously uncharacterised short open reading frames (sORFs) in the Chinese hamster genome (Figure 3a and Supplementary Data 3). sORFs are defined as ORFs predicted to produce proteins < 100 aa, termed microproteins^43^. Greater than 90% of the ORFs identified in the 5’ region of mRNAs or in transcripts previously annotated as non-coding were sORFs (Figure 3b). In this study we found 3,497 uORFs (Figure 3d) with an average length of 24 aa (Supplementary Figure 3a). AUG (46.4%) was the most prevalent start codon, followed by CUG (31%), GUG (14%) and UUG (8.4%). The average length of the ouORFs (Figure 3e) identified (n = 1,203) was 48 aa (Supplementary Figure 3b), with CUG (40.8%) the most frequent start codon, followed by AUG (27.7%), GUG (20.9%) and UUG (10.3%). The presence of uORFs in 5’ leader sequences has been shown to have a repressive effect on the main ORF in multiple species^44^, and we observed the same tendency for CHO cells in this study (see Supplementary Results).

**Figure 3:**
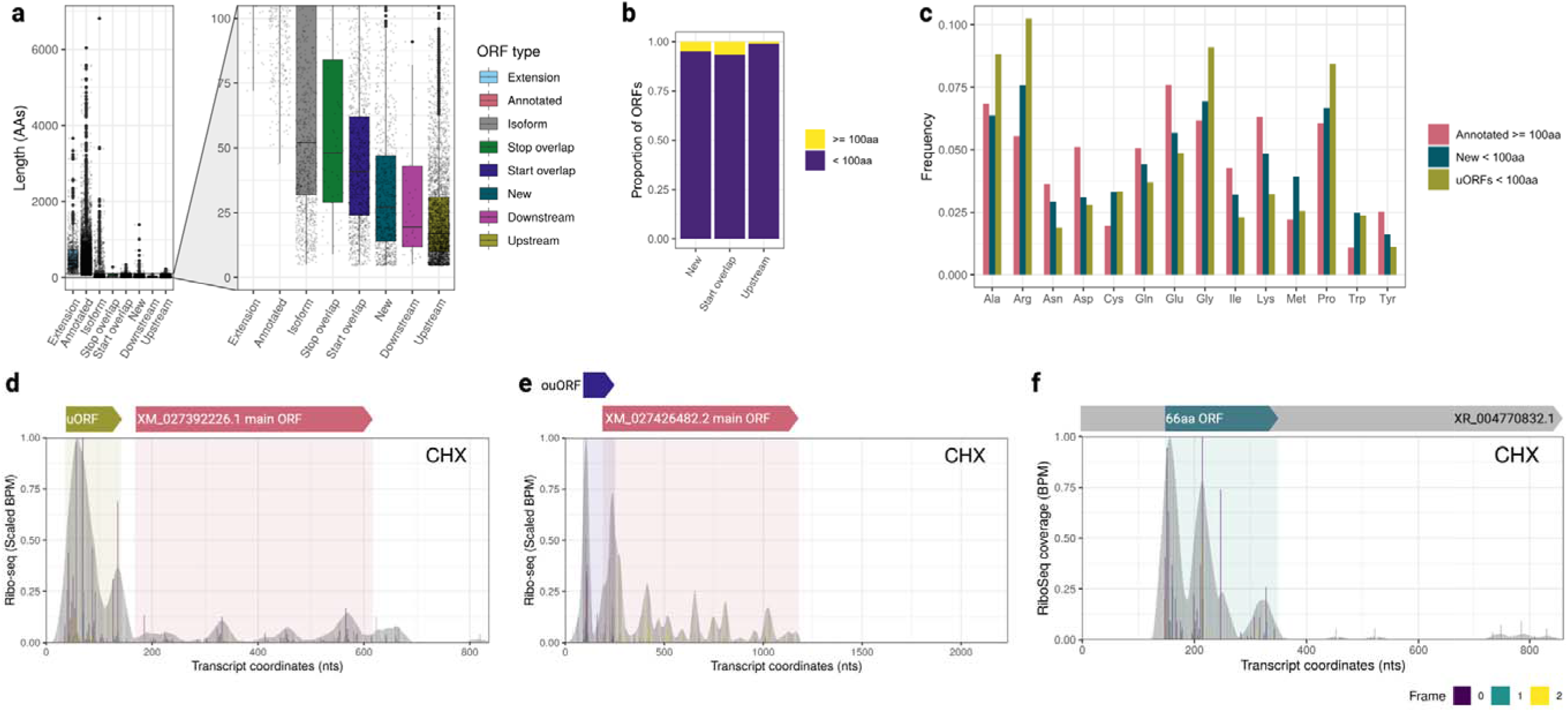
Ribosome footprint profiling uncovers thousands of short open reading frames in the Chinese hamster genome. A considerable number of previously uncharacterised ORFs identified by ORF-RATER were **(a)** predicted to be < 100 aa. In this study, we focused on short open reading frames found in the 5’ leader of protein coding transcripts (i.e., upstream ORFs and start overlapping uORFs) as well as ORFs found in non-coding RNAs where **(b)** > 90% of all identified ORFs in these classes were < 100 aa. Comparison of **(c)** the amino acid frequencies of uORFs (both uORFs and ouORFs) and ncRNA sORFs to annotated proteins, revealed differences in usage of amino acids including arginine and glycine when compared to conventional protein coding ORFs (≥ 100 aa). Examples are shown of **(d)** an uORF found in a *Ddit3* transcript, **(e)** an ouORF in *Rad51* transcript and **(f)**an sORF found in transcript designated as long non-coding RNA.

For the “New ORF” class (n = 853), the majority of ORFs were found in transcripts annotated in NCBI as non-coding (Figure 3f). The average length of “New ORFs” was 42 aa (Supplementary Figure 3c). AUG (60.4%) was the most common start codon, followed by CUG (21.9%), GUG (15.2%) and UUG (5.5%). Upstream ORFs and sORFs in the “New” ORF group, were found to have clear differences in amino acid usage, when compared to annotated proteins with ≥ 100 aa. The amino acid usage was comparable to a similar recent analysis conducted for microproteins encoded in the human genome^26^. CHO cell sORFs were found to have increased usage of arginine, glycine, and tryptophan as well as a decrease in usage of asparagine, glutamate, lysine, and aspartic acid. Alanine and proline were more prevalent in uORFs in comparison to annotated proteins and sORFs found in ncRNA, while methionine usage was more frequent in the sORFs in ncRNA (Figure 3c and Supplementary Figure 4).

### 2.4 Microproteins are a source of process-related contaminants in mAb drug products

Next, we sought to determine if the annotation of novel microproteins predicted to be encoded by sORFs increased the coverage of MS-based HCP detection. For this experiment, we utilised liquid chromatography-tandem mass spectrometry (LC-MS/MS) to analyse 4 commercial mAb drug products (pertuzumab, adalimumab, denosumab and vedolizumab) (Figure 4a). We searched the LC-MS/MS data against a protein sequence database comprised of the UniProt Chinese hamster reference proteome (n = 23,875) and selected ORFs found in this study including the sORFs predicted to encode microproteins identified from the Ribo-seq data in this study (n = 5,652) (Figure 4b). The identification of microproteins using mass spectrometry is challenging due to their size, as a lower abundance and fewer cleavage sites amenable to digestion with trypsin in comparison to canonical proteins, results in a reduction of the number of detectable peptides by MS^39^. Similar to other studies^26,45^, we considered a single peptide (≥ 4 aa) sufficient for microprotein detection while for proteins ≥ 100 aa, we employed the standard ≥ 2 peptide threshold for identification.

**Figure 4:**
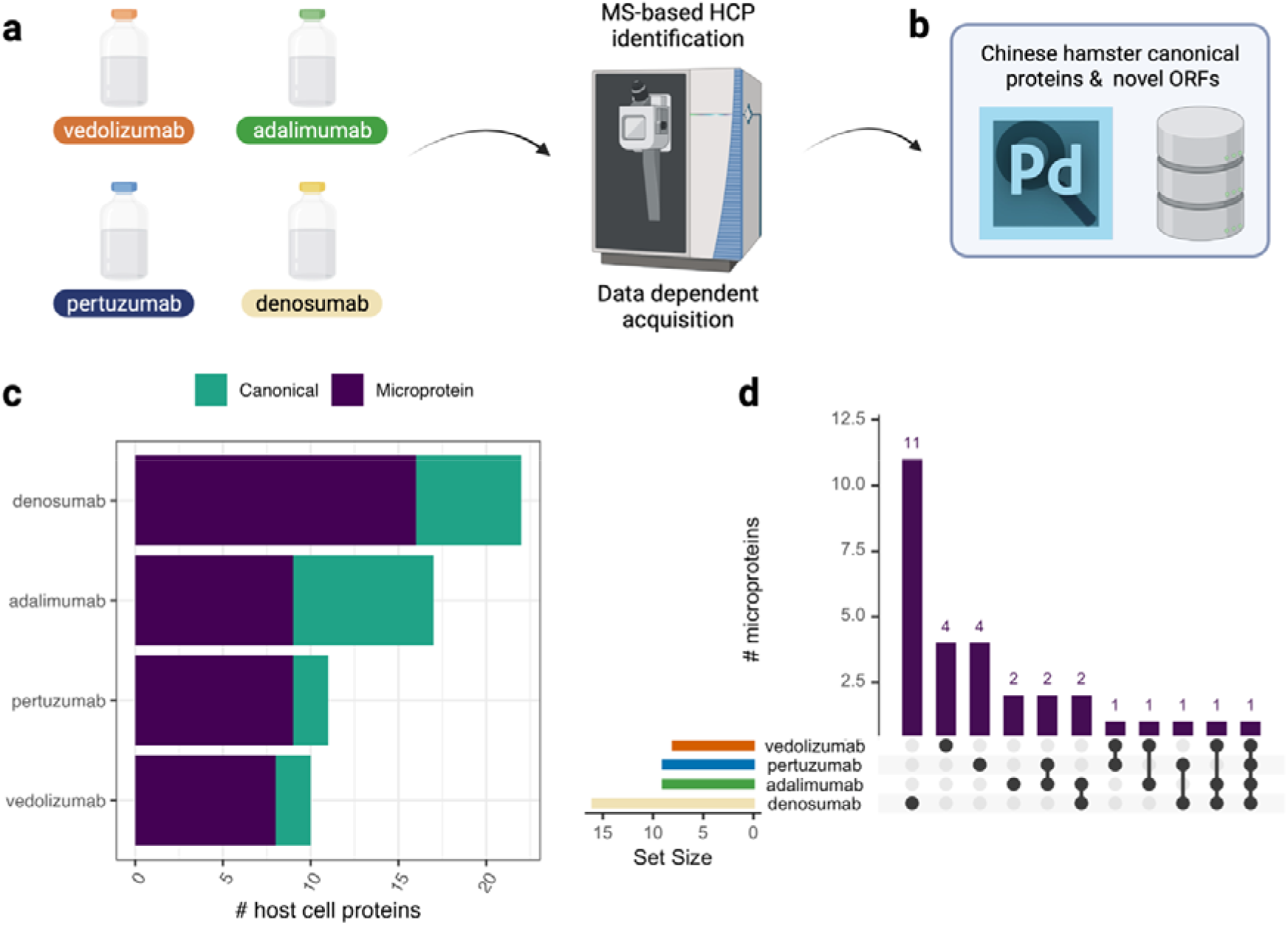
Microproteins are a new class of potential host cell impurity in mAb drug products. We conducted **(a)** LC-MS/MS based HCP analysis of 4 commercial mAb drug products (vedolizumab, adalimumab, pertuzumab, and denosumab). The resulting data were searched against a **(b)** protein database comprised of Chinese hamster proteins in UniProt and uORFs, ouORFs and non-coding RNA encoded sORFs identified using Ribo-seq. In total, we **(c)** identified 30 different microproteins across the four drug products analysed. **(d)** 10 microproteins were present in more than 1 drug product.

We detected canonical and microprotein HCPs in all 4 drug products tested in this study (Supplementary Data 4). Several previously identified canonical HCPs were found in adalimumab (e.g., cathepsin L1, S100a11^47^) and vedolizumab (e.g., Clusterin^48^). The number of microproteins found in each drug product, however, was greater than canonical proteins detected (Figure 4c). A total of 30 distinct microprotein HCPs were identified across the 4 products tested. While we find 3 microproteins with 2 or more peptides, the majority of microproteins identifications resulted from detection of a single peptide. All single-peptides identifications were from a detected peptide ≥ 10 aa. Denosumab had the largest number of individual microproteins identified (n = 16) followed by adalimumab and pertuzumab (n = 9) with vedolizumab having the lowest number identified (n = 8). Nine microproteins were found in two or more drug products, and a 36aa microprotein originating from a GUG-initiated upstream open reading frame in a *Rabl2b* transcript (XM_027396304.2) was found in all four of the drug products tested (Figure 4d).

To assess the quantities of canonical proteins microproteins present in the drug products we utilised the Hi3 HCP quantitation method^49^ (Table S4). Determination of the concentration of individual HCP using the Hi3 method requires 3 peptides to for confident identification. Three peptides were identified for 9 canonical HCPs across the 4 drug products tested and their concentration ranged from 0.98 ppm to 15.84 ppm (median = 3.40 ppm). A single 37 aa microprotein (an ouORF in a transcript of the *Cyb5d1* gene) met the 3-peptide confidence level for accurate quantitation and was found to be present in denosumab at a concentration of 2.7 ppm. Given the challenges of microprotein identification we also decided to estimate the concentration of the microproteins with fewer than 3 peptides identified (Supplementary Figure 5). Microproteins with 2 peptides identified (n = 3) ranged from 0.25 ppm to 7.90 ppm. Microproteins that were identified from a single peptide (n = 38), and therefore represented the lowest confidence estimates in terms of HCP abundance. The majority of these microproteins (n = 28) were found to be below the maximum concentration observed for canonical HCPs, while the remaining microprotein abundance estimates ranged from 19.11 to 411 ppm exceeding the canonical HCP range.

### 2.5 The translation efficiency of sORFs found in non-coding RNA genes is altered in response to a reduction of cell culture temperature

The reduction of cell culture temperature is a method used to extend the viability of some commercial cell culture processes and improve product quality^50^. A number of studies have reported that mild hypothermia can alter the abundance of canonical CHO cell HCPs^10,51–53^. Here, we wished to determine if sub-physiological temperature changed the translation efficiency of sORFs and canonical ORFs and assess if translatome analysis can provide additional valuable information on the behaviour of CHO cells. Ribo-seq enables the protein synthesis rate to be inferred by calculating the translation efficiency of each ORF. Translation efficiency is calculated by normalising the RPF occupancy by RNA abundance^16^. Significant differences in translational regulation can then be determined for each ORF following the comparison of translation efficiency between conditions^16^.

We utilised the CHX-treated Ribo-seq data for the TS and NTS sample groups, along with the matched RNA-seq data (Supplementary Figure 6). We elected to perform gene-level count-based transcriptome and translatome analysis. It is not possible to separate expression/occupancy of the uORF/ouORFs from the main ORFs using this analysis strategy. For the novel ORFs found in this study, we therefore focused only on the “New” sORFs identified in genes classified as non-coding in the reference annotation. 821 ORFs identified by ORF-RATER were found in transcripts annotated as non-coding, and 795 of these ORFs were predicted to produce a protein < 100 aa (Figure 5a). The average length of these putative microproteins found in non-coding RNA transcripts was 31 aa (Figure 5b). Most of these transcripts encoded 1 or 2 sORFs, although there were instances of up to 7 ORFs being present in a single non-coding RNA transcript (Figure 5c). To ensure compatibility with the Plastid read/RPF gene-level counting algorithm^39^ we retained only the longest sORF per non-coding transcript (collapsed to 462 genes) prior to merging with the annotated canonical ORFs for differential expression and differential translation analysis. During the read counting process, we excluded the first 5 and last 5 codons for ORFs ≥ 100 aa and the first and last codons for ORFs < 100 aa to reduce potential bias from the accumulation of ribosomes at the beginning and end of the CDS. Prior to differential expression, genes with an average < 20 counts across the 8 samples in the RNA-seq or Ribo-seq data were eliminated.

**Figure 5:**
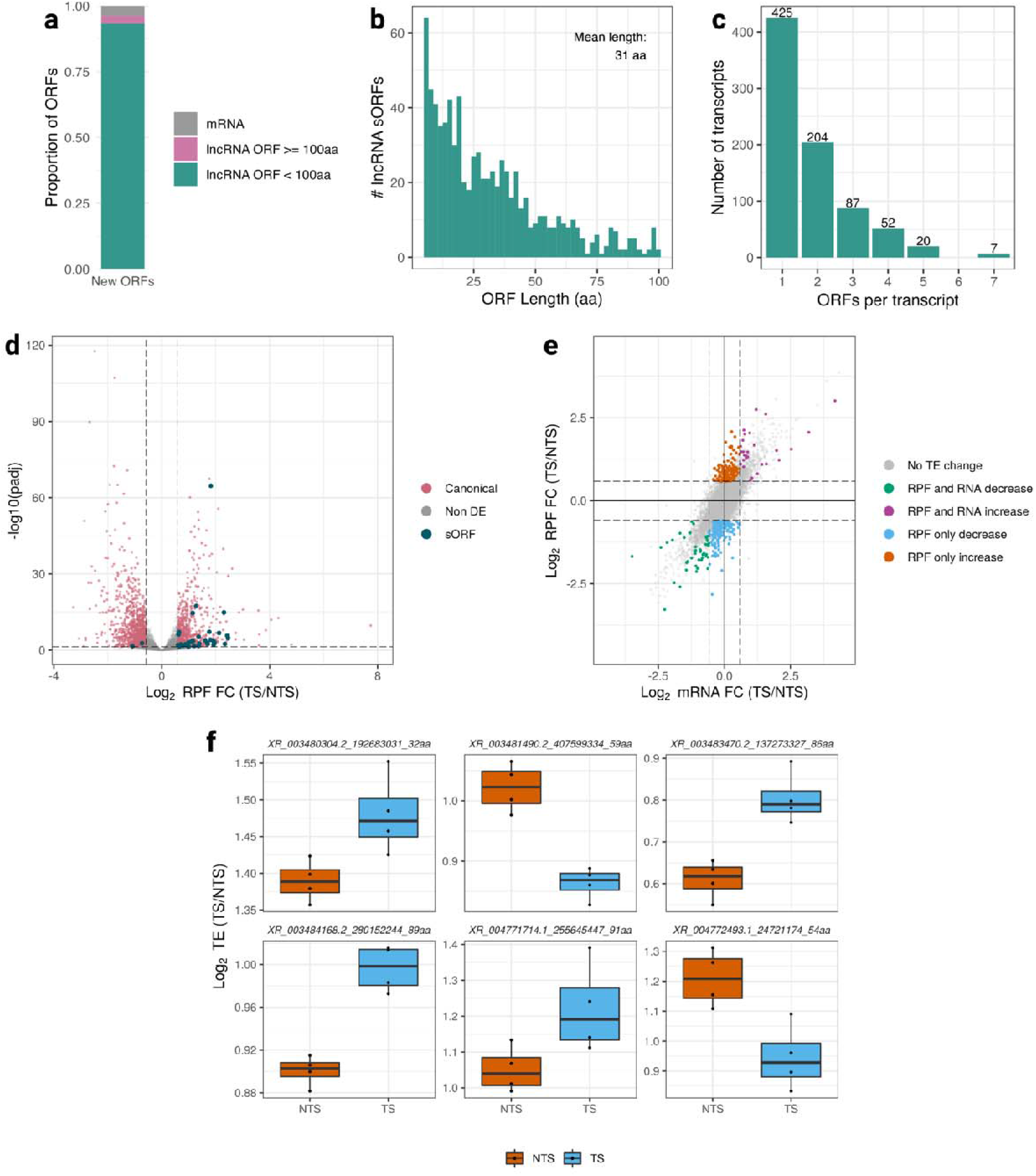
Temperature shift induces alterations in translation regulation of CHO cell canonical ORFs and sORFs. To characterise the impact of reducing cell culture temperature, we carried out differential expression and translation analysis of canonical ORFs and non-coding RNA sORFs. Of the 821 ORFs classified as “New” by ORF-RATER the majority were **(a)** in non-coding RNAs sORFs. The average length of these sORFs was **(b)** 31aa with as many as **(c)** 7 encoded by a single transcript. We found that **(d)** 1,011 ORFs (993 canonical and 18 sORFs) were significantly altered in both the RNA-seq and Ribo-seq data. Following differential translation analysis, we found that **(e)** the translational efficiency of 374 ORFs including 368 canonical ORFs and **(f)** 6 sORFs were significantly altered post-temperature shift.

We initially conducted separate analyses of the RNA-seq and Ribo-seq counts using DESeq2, to identify differences in RNA abundance (Supplementary Figure 7a, Supplementary Data S5a and S5d) and RPF occupancy (Supplementary Figure 7b, Supplementary Data S5b and S5e), as well as to determine the extent to which significant changes observed agreed for both data types. Following comparison of the TS and NTS samples, 1,880 ORFs were found to have significantly different RPF counts (1,846 canonical and 34 sORFs). 53.8% of the ORFs with significantly altered RPF density also had a significant change in RNA abundance in the same direction (Figure 5d and Supplementary Figure 7c). Gene expression and RPF occupancy differences of 18 sORFs were found to be correlated in both datasets (Supplementary Figure 8).

To identify ORFs where the translational efficiency was altered (ΔTE), DESeq2 was again utilised to assess the differences in RPF density with RNA-seq counts included as the interaction term in the model. We retained only those genes which had an average read count of 20 in both the RNA-seq and Ribo-seq datasets for this analysis. We observed significant changes in translational efficiency (≥ |1.5| fold increase or decrease in ΔTE; adjusted p-value < 0.05) for 6 sORFs and 368 canonical ORFs (Figure 5e and Figure 5f; Supplementary Data 5c and S5e). Overrepresentation analysis of canonical ORFs with altered TE, revealed that that genes in DNA repair GO biological process were enriched (FDR = 6.07 × 10^-5^) (Supplementary Data 6). Notably, a ΔTE post-temperature shift in 9 of the 26 genes overlapping with the DNA repair category, including *Brca1* (log_2_ ΔTE = -1.25 [padj = 6.3 × 10^-14^]) (Supplementary Figure 9a and 9b) was observed without a significant difference in expression of these genes in the RNA-seq data.

### 2.6 CHO cell microproteins are differentially expressed in response to mild hypothermia and between the exponential and stationary growth phases

The next step was to utilise proteomic mass spectrometry to confirm the existence of additional CHO cell microproteins. The use of mass spectrometry for direct detection also overcomes the inherent limitations of our RNA-seq and Ribo-seq analyses by enabling identification of predicted microproteins from the uORF and ouORF classes or cases when multiple microproteins are encoded by a single non-coding RNA gene. From these data, we also sought to determine if microprotein abundance was altered upon the reduction of cell culture temperature (Figure 6a). For this experiment, we again acquired cells from a non-temperature shifted control 72 hours post seeding (n = 3) and 24hr post-temperature shift (72 hours post seeding)(n = 3) as well as an additional sample at 48 hr post-temperature shift (96 hours post seeding)(n = 3) (Supplementary Figure 10a). An additional proteomics experiment was performed for a second CHO cell line to assess if microprotein abundance was altered between the exponential and stationary phases of cell growth (Figure 6b). Here, a non-mAb producing CHO-K1GS cell line was cultured for 7 days, samples were acquired for proteomics at 96h post-seeding when the cells were in exponential growth (n = 4) and at 168 hr when the cells had entered stationary phase (n = 4) (Supplementary Figure 10b). Cell lysates from both proteomics experiments were subjected to a SP3 protein clean-up procedure and tryptic digestion before LC-MS/MS-based analysis employing label-free quantification (LFQ) was performed (Figure 6c).

**Figure 6:**
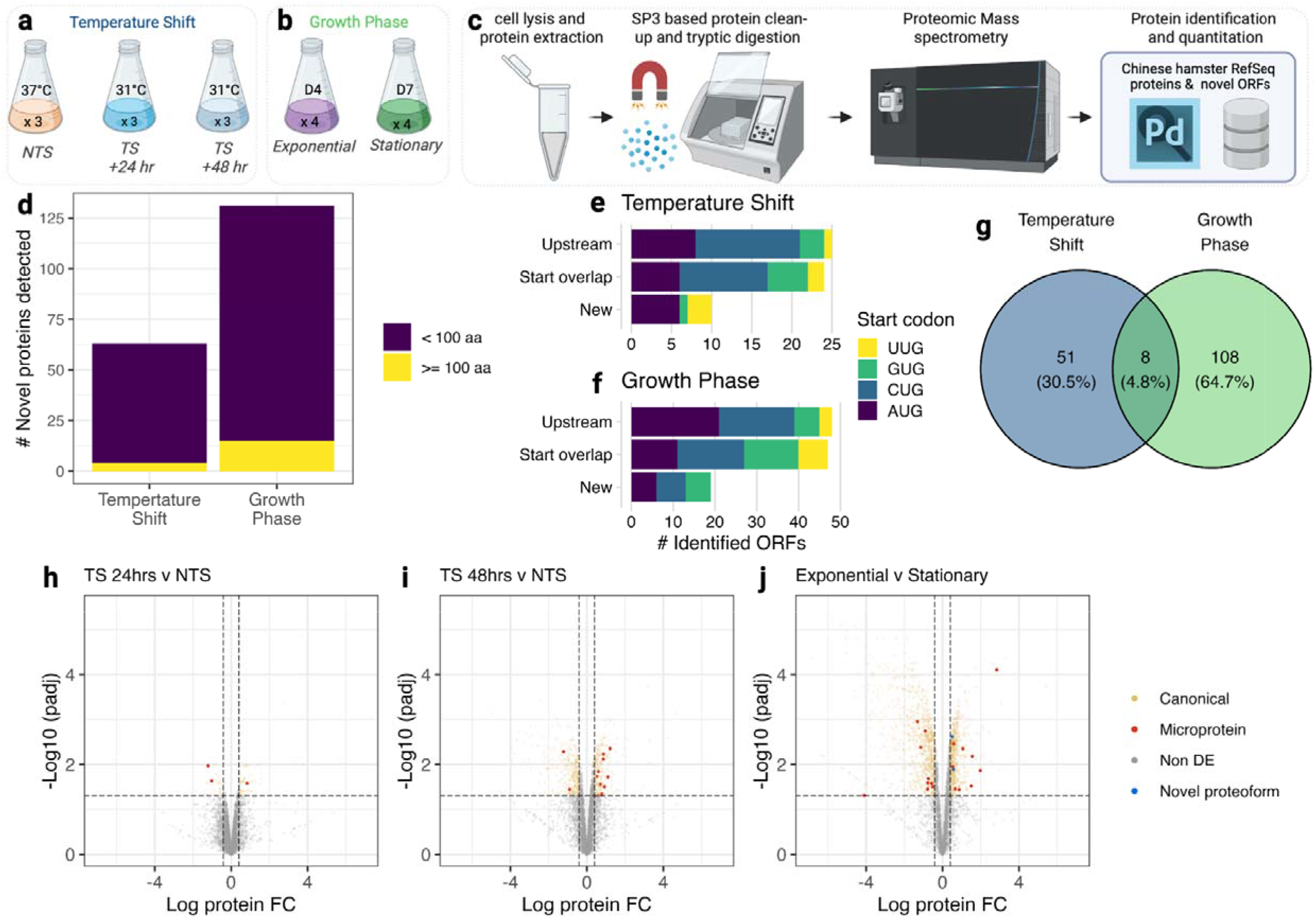
Identification of CHO cell microproteins by mass spectrometry and differential expression analysis in response to mild hypothermia and over the course of cell culture. To determine if ORFs predicted from the Ribo-seq could be identified at the protein level, we conducted LC-MS based proteomics analysis. To generate the samples for proteomics our model of **(a)** temperature shift used for Ribo-seq and RNA-seq was repeated and **(b)** cells from a non mAb-producing CHOK1GS cell line were captured during exponential growth (Day 4) and in the stationary phase (Day 7). Proteins were extracted from cell lysates and a **(c)** SP3-based protein cleanup method followed by tryptic digestion was used to prepare samples for MS analysis. The resulting data was searched against a combined database of proteins annotated for the Chinese hamster PICRH genome from RefSeq and ORF-RATER identifications using Proteome Discoverer 2.5. This analysis resulted in the identification of **(d)** 59 and 116 microproteins for the temperature shift and growth rate experiments respectively, with **(e)** 8 microproteins detected in both experiments. The microproteins identified for both experiments (**f** and **g**) originated from uORF, ouORF and non-coding RNA encoded classes. Following the comparison of protein abundances we identified differential abundances for microproteins between the non-temperature shifted control and samples acquired at **(h)** 24hrs, **(i)** 48hr post-temperature shift as well as between **(j)** the exponential and stationary phases of cell culture.

The resulting MS data from each sample was searched against the protein sequence database comprised of RefSeq protein sequences annotated for the CriGri-PICRH genome (n = 46,638) and selected ORFs including the sORFs identified from the Ribo-seq data in this study (n = 5,652). Proteins ≥ 100 aa with 2 unique peptides, and proteins < 100 aa with 1 unique peptide were considered confidently detected and retained for further analysis. For the temperature shift experiment, a total of 5,461 proteins were identified across the 9 samples analysed by mass spectrometry including 63 novel ORFs, of which 59 were classified as microproteins (Figure 6d and Supplementary Data 7) originating from the uORF (n = 25), ouORF (n = 24) and “New” ORF-RATER classes (n = 10) (Figure 6e). For the growth phase experiment, 5,084 proteins were identified including 116 microproteins (Figure 6d and Supplementary Data 8) from uORFs (n = 48), ouORFs (n = 47) and the “New” classes (n = 19) (Figure 6f). Eight microproteins were detected in both the temperature shift and growth phase experiments (Figure 6g).

The proDA algorithm^54^ was used to identify proteins significantly altered (i.e. log fold change of ≥ |1.5| and BH adjusted p-value < 0.05) between conditions for the temperature shift and growth rate experiments. Upon comparison of the 24 hr and 48 hr post-temperature shift samples to the non-temperature shifted control, 90 and 713 canonical proteins were found to be differentially expressed respectively. The abundance of 3 microproteins at 24 hr and 10 microproteins for the 48 hr post-temperature shift was observed (Figure 6h and 6i, Supplementary Data 9). In the second proteomics experiment, 1,768 canonical proteins, 17 microproteins and 2 novel ORFs ≥ 100 aa were found differentially expressed upon comparison of the D4 (exponential) and D7 (stationary) phases of cell growth (Figure 6j and Supplementary Data 10).

## 3. Discussion

Here, we present the findings of a ribosome footprint profiling experiment where both translation initiation and elongation were captured at single nucleotide resolution in CHO cells for the first time. The utilisation of harringtonine to arrest translation resulted in an enrichment of RPFs at the TIS and enabled transcriptome-wide identification of ORFs including those that started at near cognate codons. We found that the use of alternative initiation sites is widespread across the CHO cell transcriptome with ∼29% of all new ORFs identified beginning at non-AUG start codons (in agreement with TISs for the human genome present in the TISdb^55^). For previously annotated protein coding transcripts, we were able to identify 685 extended proteoforms that begin at near-cognate start codons. We also found thousands of novel sORFs predicted to encode microproteins in the 5’ leader sequence of Chinese hamster mRNAs and in ncRNA transcripts.

While the work conducted in this study has allowed us to significantly expand the annotation, we recognize that it is probable that there remain further undiscovered ORFs in the Chinese hamster genome. Future studies utilising Chinese hamster tissues as well as different CHO cell lines grown under a variety of conditions producing a range of mAb and other protein formats will enable the identification of additional ORFs. Our work is also potentially limited by the combined use of harringtonine and flash freezing, which likely led to residual elongating ribosomes and subsequent identification of potential false positive translation initiation sites. We eliminated those classes of ORFs that are liable to be affected (i.e., truncations) entirely from further analysis and conservatively assessed the remaining classes to limit false positive identifications (at the expense of potentially increasing the false negative rate). Performing future Ribo-seq experiments with different translation inhibitors such as lactimidomycin or puromycin in the future could not only enable new ORFs to be identified but also allow quantitative comparison of CHO cell translation initiation in different conditions^55,56^.

The identification of sORF-encoded CHO cell microproteins in this study has permitted the use of a more comprehensive proteomic database for mass spectrometry, resulting in an enhanced assessment of HCP impurities in four commercial mAb drug products. We identified 30 distinct host-cell microproteins using LC-MS/MS and, remarkably, the number of microproteins present was greater than canonical proteins for each drug product test. Nine microproteins were found in more than one drug product with a single microprotein found in all drug products tested. We confidently determined that a single microprotein was present in the denosumab drug product at concentration of 2.7 ppm, comparable to the median canonical HCP concentration of 3.4 ppm observed for all drug products. The majority of estimated microprotein concentrations (i.e., derived 1 or 2 peptide identifications) were found to lie within the ppm range observed for canonical HCPs. Several microprotein abundances calculated from single peptide data were found to be particularly high but it is important to note that any microprotein HCP concentration from < 3 dectected peptides reported in this study should be considered an estimate.

We wish to emphasise that we make no claims regarding any risk to the patient or impact on efficacy of the mAbs arising from the host cell microprotein impurities observed in this study. In fact, the safety and effectiveness of the more than 100 mAbs approved to date^57^, the majority of which are manufactured in CHO cells, is compelling evidence that microproteins do not cause issues, if present, in approved drug products. Nevertheless, CHO cell microproteins are a new class of host cell impurity and future studies to evaluate if, in certain circumstances, these HCPs could elicit an immune response, affect mAb stability or how they escape the purification process would be valuable for the industry. To facilitate these efforts, we have made the protein sequence database used for MS analysis freely available (<u>download</u>).

The improvement in HCP detection gained from this study is an important tool for a range of process optimisation approaches focused on limiting HCPs^58^. For instance, the HCP content in the final drug product can be influenced, in part, by the upstream process^59,60^. Here, we have shown that microprotein abundance is altered by a change in the bioreactor environment i.e., temperature. Analysis of microprotein expression during cell culture could help refine strategies that seek to modify the upstream process to control unwanted HCPs prior to harvest. Canonical HCPs present are known to vary at different stages of purification and examining their presence facilities establishment of an effective downstream process^7^. Our work enhances the ability to capture a more comprehensive snapshot of the HCP population present at each unit operation facilitating further optimisation. In recent years, several groups have reported the use of cell line engineering to reduce or knockout of problematic HCPs from CHO cell lines^11,61,62^. Through the annotation of thousands of sORFs in this study we have dramatically expanded the number of potential HCPs amenable to genome editing approaches.

We believe our work presents considerable new opportunities for continued progress in the CHO cell biology field by facilitating the study of new ORF classes as well as enhancing proteomic analyses. We have confirmed the existence of more than 170 microproteins from cell lysates using LC-MS/MS and shown that the abundance of microproteins is altered in response to temperature and over the course of cell culture. We also characterised 685 novel non-AUG initiated N-terminal extensions of canonical proteins. While AUG initiated translation is thought to result in the highest rate of protein synthesis^63^, it is also possible, as with other species^64^, that these proteoforms play important roles in bioprocess phenotypes.

Another important aspect of this study is the demonstration of the utility of analysing CHO cell translation efficiency using Ribo-seq. We have shown that the translation efficiency of canonical ORFs in genes related to DNA repair, including *Brca1*, are altered upon a decrease in cell culture temperature while no changes in the transcription of these genes was detected. We have also observed differences in RNA expression and translation efficiency of sORFs encoded in transcripts previously annotated as non-coding in response to sub-physiological cell culture temperature. Ribo-seq therefore results in a more comprehensive understanding of CHO cell behaviour than is possible with RNA-seq alone and will enable the identification of new candidates for cell line engineering studies in the future.

Our work will also be of utility to those researchers exploring approaches for the design of expression vectors for therapeutic antibody synthesis. We have shown here that ORFs in the 5’ leader of CHO cell mRNAs tend to have a repressive effect on translation of the main ORF (see Supplementary Results). The uORFs found in this study, pave the way for the use of endogenous uORFs, in a similar fashion to previous reports on synthetic uORFs^65^, to precisely control translation and/or the post-translational modification of mAbs or indeed more complicated protein formats such as bispecific antibodies^66^.

## 4. Conclusion

In conclusion, we have performed a series of Ribo-seq experiments with various translation inhibitors to examine translation elongation and, notably, translation initiation in CHO cells for the first time. Through this approach, we substantially enhanced the annotation of non-canonical ORFs in the Chinese hamster genome. We discovered novel translation events in previously annotated protein-coding genes and identified thousands of new short ORFs predicted to encode microproteins. Our findings allow improved MS-based HCP detection in mAb drug products and provide a foundation for harnessing the understanding of protein synthesis in CHO cells to further improve manufacturing process efficiency and the quality of therapeutic proteins.

## 5. Materials and Methods

### 5.1 Cell culture

#### 5.1.1 Generation of samples for Ribo-seq and RNA-seq

A mAb producing CHOK1 cell line (CHO K1 mAb) was seeded at a density of 2 × 10^5^ cells/ml in 50ml SFM-II media (Gibco, 12052098) in 8 replicate shake flasks in a Kuhner orbital shaker at 170rpm at 5% CO_2_. The cultures were grown at 37°C for 48hr post-seeding, at which point the temperature of 4 of the shake flasks was reduced to 31°C, while the remaining 4 shake flasks per experimental condition were maintained at 37°C (Figure 1A). Samples for library preparation were acquired 72 hr post-seeding. The procedure was repeated in two separate experiments, the first was used to generate Ribo-seq and matched total RNA-seq libraries from cycloheximide-treated cells (8 samples) and the second to generate Ribo-seq libraries from harringtonine-treated (8 samples) and matched no drug-treated cells.

#### 5.1.2 Generation of samples for proteomics

For proteomics analysis (Section 5.4) two experiments were conducted. The first experiment utilised an identical cell culture model for the CHO K1 mAb cell line as described in (Section 5.1.1). Here, 3 replicate samples for the non-temperature shifted control and the 24hr post temperature shift time point along with samples at 48 hr post temperature shift were acquired. The second proteomics experiment focussed on growth phases (stationary v exponential). A non-mAb CHOK1-GS cell line was seeded at a density of 2 × 10^5^ cells/ml in 30ml CD FortiCHO^TM^ medium (Gibco, cat.no. A1148301) supplemented with 4mM L-glutamine (L-Glutamine, cat.no. 25030024) in 250ml Erlenmeyer shake flasks in 8 replicates. The cultures were maintained at 37°C, 170 rpm, 5% CO_2_ and 80% humidity in a shaking incubator (Kuhner) for 4 or 7 days. On day 4 and 7 cells were counted, pelleted, and resuspended in fresh media supplemented with cycloheximide to a final concentration of 100µg/ml (Sigma, cat.no. C4859-1ml). Following a 5-minute incubation at 37°C, cells were centrifuged at 300g for 5 minutes at room temperature and the media was removed. The cell pellets were washed with ice cold PBS with cycloheximide (100µg/ml) and stored at -80=:JC until analysis.

### 5.2 Ribosome footprint profiling

#### 5.2.1 Translation Initiation sample preparation

At 72 hours post seeding, cells were treated with harringtonine (2 µg/ml) (or DMSO) for 2 minutes at 31°C or 37°C. The cultures were transferred to 50 ml tubes and following centrifugation at 1,000 rcf for 5 minutes at room temperature the media was removed, and the cells were resuspended in ice cold PBS supplemented with harringtonine or DMSO respectively. Following a 5-minute centrifugation at 1,000 rcf at 4°C, the PBS was removed, and the pellet was flash frozen in liquid nitrogen. Frozen pellets were resuspended in 400µl 1X Mammalian Polysome buffer (Illumina TruSeq Ribo Profile (mammalian) kit) prepared according to manufacturer’s guidelines. Cell lysates were incubated on ice for 10 minutes, centrifuged at 18,000 rcf for 10 minutes at 4°C to pellet cell debris and the supernatant was used for ribosome-protected fragment (RPF) isolation and library preparation.

#### 5.2.2 Translation elongation sample preparation

72 hours post seeding a total of 25 × 10^6^ cells (per replicate) were pelleted and resuspended in 20ml of fresh CHO-S-SFMII media supplemented with cycloheximide at a final concentration of 0.1 mg/mL and incubated at 37°C or 31°C for 10 min. Cells were subsequently pelleted, washed in 1 mL of ice-cold PBS containing 0.1 mg/mL of CHX, clarified and lysed. Prior to the generation of ribosomal footprints, part of the lysate was used for total RNA extraction and RNA-seq library preparation with the TruSeq Ribo Profile (mammalian) kit. PAGE Purified RPFs were used for ribosome profiling library preparation with the Illumina TruSeq Ribo Profile (mammalian) kit.

#### 5.2.3 Library preparation

To prepare RNA-seq and Ribo-seq libraries for sequencing, the TruSeq Ribo Profile (Mammalian) Kit (Illumina) was used in accordance with the manufacturer’s specifications. For Ribo-seq samples RNase treatment was performed with 10µl of TruSeq Ribo Profile Nuclease per 200µl lysate at room temperature for 45 minutes with gentle shaking. Digestion was stopped with 15µl SUPERaseIn (20U/µl) (Ambion, cat. No. AM2696). Monosomes were isolated with size exclusion chromatography using the Illustra MicroSpin S-400 HR Columns (GE life sciences, cat. no. 27514001) according to manufacturer’s instructions. Ribosomal RNA was removed with the RiboZero-Gold rRNA removal Kit (Illumina, cat. No MRZG12324). Ribosome protected fragments were size selected from a 15% denaturing urea polyacrylamide gel (PAGE) following electrophoresis (7M urea, acrylamide (19): bis-acrylamide (1)). A gel extraction step (from 15% denaturing PAGE gels) for the isolation of linker ligated ribosome protected fragments, was added to the protocol after the linker ligation reaction as in Ingolia’s protocol ^67^ for the Harringtonine and No-drug treated samples, to avoid high concentration of linker dimers contaminating the final library. Following reverse transcription, cDNA was extracted from 7.5% denaturing urea PAGE gels. PCR amplified libraries were purified from 8% PAGE gels and subsequently analysed with the Agilent High Sensitivity DNA assay (Agilent, Bioanalyzer).

#### 5.2.4 Sequencing

The libraries for translation initiation and elongation analyses were sequenced on an Illumina NextSeq configured to yield 75bp and 50bp single end reads respectively.

### 5.3 Ribo-seq and RNA-seq data analysis

#### 5.3.1 Pre-processing

Adapter sequences were trimmed from the Ribo-seq and RNA-seq datasets using Cutadapt v1.18^68^, and Trimmomatic v0.36^69^ was used to remove low quality bases. To remove contaminants from the Ribo-seq data Chinese hamster rRNA, tRNA and snoRNA sequences were downloaded from v18 of the RNAcentral v18 database^38^ and an individual STAR v2.7.8a^37^ index was built for each type of RNA. The Ribo-seq reads were aligned against each index using the following parameters: -- seedSearchStartLmaxOverLread .5 --outFilterMultimapNmax 1000 --outFilterMismatchNmax 2. Reads that mapped to rRNA, tRNA or snoRNA were discarded.

#### 5.3.2 Read alignment

The pre-processed Ribo-seq and RNA-seq data were aligned to the NCBI CriGri-PICRH 1.0 genome and transcriptome (GCA_003668045.2) ^14^ with STAR v2.7.8a using the following parameters: -- outFilterMismatchNmax 2 --outFilterMultimapNmax 1 --outFilterMatchNmin 16 --aligEndsType EndToEnd.

#### 5.3.3 Ribo-seq P-site offset identification and selection of RPFs

The P-site offset (the number of nucleotides between the 5’ end of a Ribo-seq read and the P-site of the ribosome footprint that was captured) was determined using Plastid v0.4.8^39^ by first defining the genomic region around annotated Chinese hamster CDS using the metagene generate programme with default settings. The Plastid P-site tool was then used to assess the P-site for different read lengths around the expected mammalian RPF size (27-32nt) for CDSs that had at least 10 mapped reads to the start region. Following the determination of P-site offsets for each read length for the CHX, HARR and ND Ribo-seq data, only those read lengths where ≥ 60% of the reads were found to have the expected triplet periodicity with a P-site offset of 12 were retained for further analysis.

#### 5.3.4 ORF identification

The 8 replicates from each Ribo-seq type were merged to increase sensitivity before the ORF-RATER pipeline ^40^ was used to identify ORFs in the Chinese hamster genome. Annotated pseudogenes were removed from the reference with only those transcripts with a minimum of 64 mapped RPFs from the CHX and ND Ribo-seq data were considered for ORF identification. The ORF search was limited to NUG codons with only the HARR Ribo-seq data used to identify the translation initiation sites, while the CHX and ND RPFs were used to assess translation at putative ORFs. Identified ORFs with an ORF-RATER score ≥ 0.5^41,42^ and with a length ≥ 5aa were retained. Visualisation of transcripts with novel Chinese hamster ORFs was accomplished using deeptools bamCoverage^70^. Where one or more Ribo-seq type was displayed on the same figure, the bins per million (BPM) value was scaled between 0 and 1.

#### 5.3.5 Transcript-level quantitation

The RNA abundance and RPF density in reads per kilobase mapped (RPKM) of annotated and novel ORFs was determined for each CHX-treated Ribo-seq replicate from the NTS and TS samples using the Plastid cs programme^39^. Reads and RPFs aligning to the first 5 or last 15 codons of each CDS were eliminated for ORFs ≥ 100 aa while for those ORFs < 100 aa the first and last codon counts were excluded^26^. The translation efficiency for CDS regions was calculated by dividing the RPF RPKM value by that of the RPKM of the matched RNA-seq sample.

#### 5.3.6 Gene-level differential expression and differential translation analysis

To conduct gene-level differential translation analysis the reference protein coding annotation was merged with selected ORFs found to be encoded by non-coding RNAs. Prior to counting Plastid cs generate was used to collapse transcripts that shared exons, remove regions comprised of more than 1 same-strand gene and create position groups corresponding to exons, CDS, 5’ leader and 3’UTR. An identical codon masking procedure to transcripts was also utilised for gene level analyses. The counts corresponding to CDS regions were analysed by DESeq2^71^ to identify differences between the RNA-seq and RPF counts for the TS and NTS groups. For differential translation analysis, the RNA-seq data was used as an interaction term within the DESeq2 model to enable the identification of changes in RPF density independently of RNA abundance. For all analyses, an absolute fold change ≥ 1.5 and BH adjusted p-value < 0.05 were considered significant. Gene level coverage was determined using deepTools bamCoverage and the wiggleplotr R-package v1.16.0^72^ was used to display the track and corresponding gene model.

#### 5.3.7 Enrichment analysis

The overrepresentation of gene ontology (GO) biological processes in differentially expressed and/or differentially translated genes were identified with the R WebGestaltR package^73^. Where no gene symbol was available the Chinese hamster gene name was mapped to the NCBI Mus musculus GRCm39 annotation, and the corresponding mouse gene symbol was used. GO biological processes with a BH adjusted p-value of < 0.05 were considered significant.

### 5.4 CHO cell proteomics

#### 5.4.1 Sample preparation for reversed phase liquid chromatography-tandem mass spectrometry (RPLC-MS/MS)

Samples obtained from the temperature shift and growth phase experiments (Section 5.1.2) were prepared for proteomics using a semi-automated version of the SP3 protocol^74^. Briefly CHO cells were pelleted via centrifugation at 300 × g for 5 mins. Following a wash step with 1 × PBS, cells were lysed using 1 × RIPA buffer (Cell Signalling Technology, Dublin, Ireland) containing 1 × protease inhibitor (cOmplete™, Mini, EDTA-free Protease Inhibitor Cocktail, Sigma, Wicklow, Ireland) followed by sonication. After removing cell debris *via* centrifugation, the protein concentration was determined and an aliquot of the sample containing 50 µg of protein was used for tryptic digestion as described previously^75^. Following digestion, magnetic beads were removed, and samples were acidified by adding 0.1% (v/v) formic acid before LC-MS analysis (Section 5.4.2 and Section 5.4.3). Note: An identical sample preparation procedure was carried out for HCP analysis of mAb drug products (Section 5.5.2).

#### 5.4.2 Temperature shift experiment RPLC-MS/MS analysis

CHO K1 mAb cell lysates obtained after reducing cell culture temperature were analysed using an Orbitrap Exploris™ 480 mass spectrometer (Thermo Fisher Scientific, Bremen, Germany) online hyphenated to an UltiMate™ 3000 RSLCnano system using an EASY-Spray™ source. 1 µg per sample was loaded onto a C18 Nano-Trap column followed by separation using an EASY-Spray Acclaim PepMap 100, 75 µm × 50 cm column maintained at 45.0°C at a flow rate of 250.0 nL/min. Separation was achieved using a gradient of (A) 0.10% (v/v) formic acid in water and (B) 0.10% (v/v) formic acid in acetonitrile. Gradient conditions were as follows: 2-25% B in 120 min, followed by another increase to 45% B in 30 min. The separation was followed by 2 wash steps at 80% B for 5 min and the column was re-equilibrated at 5% B for 15 min.

MS detection was carried out in centroid positive ion mode. First, full scans were acquired at a resolution setting of 60,000 (at *m/z* 200) with a scan range or *m/z* 200-2,000. The normalised automatic gain control (AGC) target was set to 100% with a maximum IT of 50 ms. 20 most abundant precursor ions were selected for HCD fragmentation using a normalized collision energy of 28%. For isolation of precursor ions, an isolation window of 1.2 *m/z* and an intensity threshold of 5.0e3 was used. Fragment scans were acquired using a resolution setting of 15,000 (at *m/z* 200) with an AGC target of 50% and a maximum IT of 70 ms. Unassigned charge states and charge states >7, were excluded from fragmentation. A dynamic exclusion was used for 45 seconds with a tolerance of ± 5 ppm.

#### 5.4.3 Growth phase experiment RPLC-MS/MS analysis

Mass spectrometric analysis was performed using an Orbitrap Eclipse™ Tribid™ mass spectrometer (Thermo Fisher Scientific, Bremen, Germany) coupled to an UltiMate™ 3000 RSLCnano system by means of an EASY-Spray™ source (Thermo Fisher Scientific, Germering, Germany). 2 µg per sample were loaded onto a C18 Nano-Trap Column followed by separation using an EASY-Spray Acclaim PepMap 100, 75 µm × 50 cm column (Thermo Fisher Scientific, Sunnyvale, CA, United States) maintained at 45.0°C at a flow rate of 250.0 nL/min. Separation was achieved using a gradient of (A) 0.10% (v/v) formic acid in water and (B) 0.10% (v/v) formic acid in acetonitrile (LC-MS optima, Fisher Scientific). Gradient conditions were as follows: 5% B for 5 min, followed by a linear gradient of 5-25% in 95 min, followed by another increase to 35% B in 20 min. The separation was followed by 2 wash steps at 90% B for 5 min and the column was re-equilibrated at 5% B for 15 min.

MS analysis was performed in positive ion mode. Full scans were acquired in the Orbitrap at a resolution setting of 120,000 (at m/z 200) with a scan range of m/z 200-2,000 using a normalized AGC target of 100% and an automatic maximum injection time in centroid mode. Using a 3 s cycle time, ions were selected for HCD fragmentation using a collision energy setting of 28%. Fragment scans were acquired in the Orbitrap using a resolution setting of 30,000 (at m/z 200). The AGC target was set to 200%. For isolation of precursor ions, an isolation window of 1.2 m/z was used. An intensity threshold of 5.0e4 was applied while unassigned charge states as well as charges >6 were excluded. The dynamic exclusion time was set to 60 s with ± 5 ppm tolerance.

#### 5.4.4 Proteome discoverer data analysis

Analysis of acquired raw data was performed in Proteome Discoverer version 2.5 (Thermo Fischer Scientific). Each dataset was searched against a protein sequence database comprised of RefSeq Protein sequences for the CriGri-PICRH genome (n = 46,638) and the novel ORFs (n = 5,652).

#### 5.4.5 Protein detection and quantitation in cell lysate samples

For protein identification as well as label-free quantitation (LFQ) in cell lysate samples, two Sequest HT searches were performed using fixed value PSM validator (#1 and #2) and an extra Sequest HT (#3) search was conducted using Percolator. Detailed settings of the database search can be found in Supplementary Data 11a.

#### 5.4.6 Identification of differentially expressed proteins

LFQ data was log2 transformed, normalised on total peptide amount and the proDA algorithm^54^ was then utilised to fit a probabilistic drop out model to these data prior to differential expression analysis. Proteins with a ≥ |1.5| fold change, adjusted p-value < 0.05 and detected in at least 3 or 4 samples for the temperature shift and growth phase experiment respectively were considered differentially expressed.

### 5.5 Host cell protein analysis

#### 5.5.1 RPLC-MS/MS analysis of drug products

Following tryptic digestion as described above (Section 5.4.1), HCP analysis of the pertuzumab, adalimumab, denosumab and vedolizumab commercial drug products (Evidentic, Berlin, Germany) was performed using a Orbitrap Exploris™ 480 mass spectrometer coupled to a Vanquish™ Neo UHPLC system (Thermo Fisher Scientific, Germering, Germany) using an EASY-Spray™ source. LC-MS/MS analysis was performed in two steps: First, 100 ng/sample were analysed to generate an exclusion list containing drug product derived peptides. Second, 3 µg were analysed in triplicate using the corresponding exclusion lists to facilitate HCP detection. For both experiments, similar LC-MS parameters were used as outlined below. For quantitation, Hi3 E. coli (Waters, Milford, MA, USA) standard was added to reach a final concentration of 50 fmol/µg of protein injected.

Using the pressure driven injection mode, either 100 ng or 3 µg of sample were loaded onto a C18 Nano-Trap column followed by separation using a 50 cm × 75 µm EASY-Spray™ PepMap™ Neo UHPLC column (Thermo Fisher Scientific, Sunnyvale, CA, USA). For separation, a linear gradient of 2-25% B (0.1% formic acid (v/v) in acetonitrile) over 60 min followed by another increase to 45% B in 30 min was used. Separation was followed by 2 wash steps at 80% B prior to column re-equilibration using 3 column volumes. The flow rate was 250nl/min and column temperature were maintained at 45.0°C.

Data dependent MS/MS analysis was performed in positive ion mode. First, full scans were acquired covering a scan range of *m/z* 200-2000 using a resolution setting of 60,000 (at *m/z* 200). The AGC target was set to 100% with a maximum IT of 50 ms. 20 most abundant ions were selected for fragmentation using a HCD collision energy of 28%. An isolation window of 1.2 *m/z* was used. The AGC target for MS2 scans acquired using a resolution setting of 15,000 (at *m/z* 200) was set to 50% with a maximum IT of 70 ms. Only charge states of +2 to +7 were used for fragmentation. Additionally, an intensity threshold of 5.0e3 was applied. Dynamic exclusion was used for 45 s with a ± 5 ppm tolerance. For targeted mass exclusion of drug product derived peptides, a retention time window of 5 min was allowed as well as a tolerance of 10 ppm.

#### 5.5.2 Detection and quantitation of HCPs in drug product samples

For detection and Hi3 quantitation of HCPs in drug product samples, two Sequest HT searches were performed using Fixed value PSM validator against a database comprised of proteins in the UniProt Chinese hamster reference proteome (n = 3,875) and the sORF containing proteoform classes identified from the Ribo-seq data in this study (n = 5,652). Detailed settings of the database search can be found in Supplementary Data 11b. Prior to further data analysis, results were filtered for contaminants and only proteins with a sequence coverage greater than 1% and Sequest HT scores ≠ 0.0 were considered.

### 5.6 Data availability

The Ribo-seq and RNA-seq data from the harringtonine, cycloheximide and no-drug treated cells have been deposited in the Sequence Read Archive (SRA) with accession code <u>PRJNA778050</u>. The code required to reproduce the results presented in this manuscript is available at https://github.com/clarke-lab/CHO_cell_microprotein_analysis. The proteomics data have been deposited to the ProteomeXchange Consortium via the PRIDE partner repository with the dataset identifier <u>PXD030186</u>. The protein sequence database used for HCP and proteomics analysis is available at https://doi.org/10.5281/zenodo.5801357.

## Supporting information

Description of Supplementary Data

Supplementary Results

## Abbreviations

BH: Benjamini-Hochberg
CDS: coding sequence
CHO: Chinese hamster ovary
CHX: cycloheximide
EST: Expressed sequence tag
Harr: Harringtonine
HCP: host cell protein
mAb: monoclonal antibody
LC-MS/MS: liquid chromatography-tandem mass spectrometry
MS: mass spectrometry
ELISA: Enzyme-linked immunosorbent assays
NGS: Next generation sequencing
NTS: non-temperature shifted
ORF: open reading frame
ouORF: overlapping upstream ORF
PAGE: polyacrylamide gel
Ribosome footprint profiling: Ribo-seq
RPF: Ribosome protected fragment
RPKM: Reads per kilobase mapped
sORF: short open reading frame
TIS: Translation Initiation Site
TS: temperature shifted
TE: Translational efficiency
uORF: upstream open reading frame
UTR: untranslated region
BPM: Bins per million
AGC: Automatic Gain Control
GO: Gene Ontology
LFQ: Label Free Quantification
DDA: Data Dependent Acquisition
IT: Injection Time
RPLC-MS/MS: reversed phase liquid chromatography-tandem mass spectrometry

## Author Contributions

IT and CC conceived the study and designed experiments; Cell culture and Ribo-seq were carried by IT, MCh and PK. Ribo-seq data analysis was performed by MCR and CC. CT, FG, LS, and JB performed the proteomics analysis. MCR, IT, LZ, MCl, BLK, NB, JB and CC wrote the manuscript. All authors reviewed the paper.

## Acknowledgements

The authors gratefully acknowledge funding from Science Foundation Ireland (grant references: 15/CDA/3259 and 13/SIRG/2084). Figures were created with BioRender.com.

## References

1. Walsh, G. & Walsh, E. Biopharmaceutical benchmarks 2022. Nat. Biotechnol. 40, 1722–1760 (2022).

2. Hanania, N. A. et al. Lebrikizumab in moderate-to-severe asthma: pooled data from two randomised placebo-controlled studies. Thorax 70, 748–756 (2015).

3. Li, X. et al. Identification and characterization of a residual host cell protein hexosaminidase B associated with N-glycan degradation during the stability study of a therapeutic recombinant monoclonal antibody product. Biotechnol. Prog. 37, e3128 (2021).

4. Luo, H. et al. Cathepsin L Causes Proteolytic Cleavage of Chinese-Hamster-Ovary Cell Expressed Proteins During Processing and Storage: Identification, Characterization, and Mitigation. Biotechnol. Prog. 35, e2732 (2019).

5. Bracewell, D. G., Francis, R. & Smales, C. M. The future of host cell protein (HCP) identification during process development and manufacturing linked to a risk-based management for their control. Biotechnol. Bioeng. 112, 1727–1737 (2015).

6. Zhu-Shimoni, J. et al. Host cell protein testing by ELISAs and the use of orthogonal methods. Biotechnol. Bioeng. 111, 2367–2379 (2014).

7. Pilely, K. et al. Monitoring process-related impurities in biologics–host cell protein analysis. Anal. Bioanal. Chem. 414, 747–758 (2022).

8. Henry, S. M., Sutlief, E., Salas-Solano, O. & Valliere-Douglass, J. ELISA reagent coverage evaluation by affinity purification tandem mass spectrometry. mAbs 9, 1065–1075 (2017).

9. Huang, Y., Molden, R., Hu, M., Qiu, H. & Li, N. Toward unbiased identification and comparative quantification of host cell protein impurities by automated iterative LC–MS/MS (HCP-AIMS) for therapeutic protein development. J. Pharm. Biomed. Anal. 200, 114069 (2021).

10. Goey, C. H., Bell, D. & Kontoravdi, C. Mild hypothermic culture conditions affect residual host cell protein composition post-Protein A chromatography. mAbs 10, 476–487 (2018).

11. Chiu, J. et al. Knockout of a difficult-to-remove CHO host cell protein, lipoprotein lipase, for improved polysorbate stability in monoclonal antibody formulations. Biotechnol. Bioeng. 114, 1006–1015 (2017).

12. Xu, X. et al. The genomic sequence of the Chinese hamster ovary (CHO)-K1 cell line. Nat. Biotechnol. 29, 735–741 (2011).

13. Meleady, P. et al. Utilization and evaluation of CHO-specific sequence databases for mass spectrometry based proteomics. Biotechnol. Bioeng. 109, 1386–1394 (2012).

14. Hilliard, W., MacDonald, M. L. & Lee, K. H. Chromosome-scale scaffolds for the Chinese hamster reference genome assembly to facilitate the study of the CHO epigenome. Biotechnol. Bioeng. 117, 2331–2339 (2020).

15. Li, S. et al. Proteogenomic annotation of the Chinese hamster reveals extensive novel translation events and endogenous retroviral elements. J. Proteome Res. 18, 2433–2445 (2019).

16. Ingolia, N. T., Ghaemmaghami, S., Newman, J. R. S. & Weissman, J. S. Genome-Wide Analysis in Vivo of Translation with Nucleotide Resolution Using Ribosome Profiling. Science 324, 218– 223 (2009).

17. Wright, B. W., Yi, Z., Weissman, J. S. & Chen, J. The dark proteome: translation from noncanonical open reading frames. Trends Cell Biol. S0962–8924(21)00226–9 (2021) doi:10.1016/j.tcb.2021.10.010.

18. Mudge, J. M. et al. Standardized annotation of translated open reading frames. Nat. Biotechnol. 40, 994–999 (2022).

19. Ivanov, I. P., Firth, A. E., Michel, A. M., Atkins, J. F. & Baranov, P. V. Identification of evolutionarily conserved non-AUG-initiated N-terminal extensions in human coding sequences. Nucleic Acids Res. 39, 4220–4234 (2011).

20. Ji, Z., Song, R., Regev, A. & Struhl, K. Many lncRNAs, 5’UTRs, and pseudogenes are translated and some are likely to express functional proteins. eLife 4, e08890 (2015).

21. Zhang, H. et al. Determinants of genome-wide distribution and evolution of uORFs in eukaryotes. Nat. Commun. 12, 1076 (2021).

22. Aspden, J. L. et al. Extensive translation of small Open Reading Frames revealed by Poly-Ribo-Seq. eLife 3, e03528 (2014).

23. Bazzini, A. A. et al. Identification of small ORFs in vertebrates using ribosome footprinting and evolutionary conservation. EMBO J. 33, 981–993 (2014).

24. Ingolia, N. T., Lareau, L. F. & Weissman, J. S. Ribosome profiling of mouse embryonic stem cells reveals the complexity and dynamics of mammalian proteomes. Cell 147, 789–802 (2011).

25. Chen, J. et al. Pervasive functional translation of noncanonical human open reading frames. Science 367, 1140–1146 (2020).

26. Martinez, T. F. et al. Accurate annotation of human protein-coding small open reading frames. Nat. Chem. Biol. 16, 458–468 (2020).

27. Zhang, S. et al. Mitochondrial peptide BRAWNIN is essential for vertebrate respiratory complex III assembly. Nat. Commun. 11, 1312 (2020).

28. Rathore, A. et al. MIEF1 Microprotein Regulates Mitochondrial Translation. Biochemistry 57, 5564–5575 (2018).

29. Lee, C. et al. The mitochondrial-derived peptide MOTS-c promotes metabolic homeostasis and reduces obesity and insulin resistance. Cell Metab. 21, 443–454 (2015).

30. Slavoff, S. A., Heo, J., Budnik, B. A., Hanakahi, L. A. & Saghatelian, A. A human short open reading frame (sORF)-encoded polypeptide that stimulates DNA end joining. J. Biol. Chem. 289, 10950–10957 (2014).

31. Koh, M. et al. A short ORF-encoded transcriptional regulator. Proc. Natl. Acad. Sci. U. S. A. 118, e2021943118 (2021).

32. Kuo, C.-C. et al. The emerging role of systems biology for engineering protein production in CHO cells. Curr. Opin. Biotechnol. 51, 64–69 (2018).

33. Donaldson, J., Kleinjan, D.-J. & Rosser, S. Synthetic biology approaches for dynamic CHO cell engineering. Curr. Opin. Biotechnol. 78, 102806 (2022).

34. Kallehauge, T. B. et al. Ribosome profiling-guided depletion of an mRNA increases cell growth rate and protein secretion. Sci. Rep. 7, 40388 (2017).

35. Masterton, R. J. & Smales, C. M. The impact of process temperature on mammalian cell lines and the implications for the production of recombinant proteins in CHO cells. Pharm. Bioprocess. 2, 49–61 (2014).

36. Tzani, I. et al. Subphysiological temperature induces pervasive alternative splicing in Chinese hamster ovary cells. Biotechnol. Bioeng. 117, 2489–2503 (2020).

37. Dobin, A. et al. STAR: ultrafast universal RNA-seq aligner. Bioinformatics 29, 15–21 (2013).

38. The RNAcentral Consortium. RNAcentral: a hub of information for non-coding RNA sequences. Nucleic Acids Res. 47, D221–D229 (2019).

39. Dunn, J. G. & Weissman, J. S. Plastid: nucleotide-resolution analysis of next-generation sequencing and genomics data. BMC Genomics 17, 958 (2016).

40. Fields, A. P. et al. A Regression-Based Analysis of Ribosome-Profiling Data Reveals a Conserved Complexity to Mammalian Translation. Mol. Cell 60, 816–827 (2015).

41. Eisenberg, A. R. et al. Translation Initiation Site Profiling Reveals Widespread Synthesis of Non-AUG-Initiated Protein Isoforms in Yeast. Cell Syst. 11, 145–160.e5 (2020).

42. Finkel, Y. et al. The coding capacity of SARS-CoV-2. Nature 589, 125–130 (2021).

43. Olexiouk, V., Van Criekinge, W. & Menschaert, G. An update on sORFs.org: a repository of small ORFs identified by ribosome profiling. Nucleic Acids Res. 46, D497–D502 (2018).

44. Chew, G.-L., Pauli, A. & Schier, A. F. Conservation of uORF repressiveness and sequence features in mouse, human and zebrafish. Nat. Commun. 7, 11663 (2016).

45. Zhu, Y. et al. Discovery of coding regions in the human genome by integrated proteogenomics analysis workflow. Nat. Commun. 9, 903 (2018).

46. Ahrens, C. H., Wade, J. T., Champion, M. M. & Langer, J. D. A Practical Guide to Small Protein Discovery and Characterization Using Mass Spectrometry. J. Bacteriol. 204, e00353–21 (2022).

47. Füssl, F. et al. Comprehensive characterisation of the heterogeneity of adalimumab via charge variant analysis hyphenated on-line to native high resolution Orbitrap mass spectrometry. mAbs 11, 116–128 (2019).

48. Zhang, Q. et al. Comprehensive tracking of host cell proteins during monoclonal antibody purifications using mass spectrometry. mAbs 6, 659–670 (2014).

49. Strasser, L. et al. Detection and quantitation of host cell proteins in monoclonal antibody drug products using automated sample preparation and data-independent acquisition LC-MS/MS. J. Pharm. Anal. 11, 726–731 (2021).

50. Masterton, R. J. & Smales, C. M. The impact of process temperature on mammalian cell lines and the implications for the production of recombinant proteins in CHO cells. Pharm. Bioprocess. 2, 49–61 (2014).

51. Goey, C. H., Tsang, J. M. H., Bell, D. & Kontoravdi, C. Cascading effect in bioprocessing-The impact of mild hypothermia on CHO cell behavior and host cell protein composition. Biotechnol. Bioeng. 114, 2771–2781 (2017).

52. Jin, M., Szapiel, N., Zhang, J., Hickey, J. & Ghose, S. Profiling of host cell proteins by two-dimensional difference gel electrophoresis (2D-DIGE): Implications for downstream process development. Biotechnol. Bioeng. 105, 306–316 (2010).

53. Tait, A. S., Tarrant, R. D. R., Velez-Suberbie, M. L., Spencer, D. I. R. & Bracewell, D. G. Differential Response in Downstream Processing of CHO Cells Grown Under Mild Hypothermic Conditions. Biotechnol. Prog. 29, 688–696 (2013).

54. Ahlmann-Eltze, C. & Anders, S. proDA: Probabilistic dropout analysis for identifying differentially abundant proteins in label-free mass spectrometry. Biorxiv 661496 (2020).

55. Lee, S. et al. Global mapping of translation initiation sites in mammalian cells at single-nucleotide resolution. Proc. Natl. Acad. Sci. U. S. A. 109, E2424–2432 (2012).

56. Zhang, P. et al. Genome-wide identification and differential analysis of translational initiation. Nat. Commun. 8, 1749 (2017).

57. Mullard, A. FDA approves 100th monoclonal antibody product. Nat. Rev. Drug Discov. 20, 491– 495 (2021).

58. Tuameh, A., Harding, S. E. & Darton, N. J. Methods for addressing host cell protein impurities in biopharmaceutical product development. Biotechnol. J. 18, e2200115 (2023).

59. Wilson, L. J., Lewis, W., Kucia-Tran, R. & Bracewell, D. G. Identification of upstream culture conditions and harvest time parameters that affect host cell protein clearance. Biotechnol. Prog. 35, e2805 (2019).

60. Hogwood, C. E., Bracewell, D. G. & Smales, C. M. Measurement and control of host cell proteins (HCPs) in CHO cell bioprocesses. Curr. Opin. Biotechnol. 30, 153–160 (2014).

61. Fukuda, N., Senga, Y. & Honda, S. Anxa2- and Ctsd-knockout CHO cell lines to diminish the risk of contamination with host cell proteins. Biotechnol. Prog. 35, e2820 (2019).

62. Kol, S. et al. Multiplex secretome engineering enhances recombinant protein production and purity. Nat. Commun. 11, 1908 (2020).

63. Kearse, M. G. & Wilusz, J. E. Non-AUG translation: a new start for protein synthesis in eukaryotes. Genes Dev. 31, 1717–1731 (2017).

64. Liang, H. et al. PTENα, a PTEN isoform translated through alternative initiation, regulates mitochondrial function and energy metabolism. Cell Metab. 19, 836–848 (2014).

65. Ferreira, J. P., Overton, K. W. & Wang, C. L. Tuning gene expression with synthetic upstream open reading frames. Proc. Natl. Acad. Sci. U. S. A. 110, 11284–11289 (2013).

66. Ong, H. K., Nguyen, N. T. B., Bi, J. & Yang, Y. Vector design for enhancing expression level and assembly of knob-into-hole based FabscFv-Fc bispecific antibodies in CHO cells. Antib. Ther. 5, 288 (2022).

67. Ingolia, N. T., Brar, G. A., Rouskin, S., McGeachy, A. M. & Weissman, J. S. The ribosome profiling strategy for monitoring translation in vivo by deep sequencing of ribosome-protected mRNA fragments. Nat. Protoc. 7, 1534–1550 (2012).

68. Martin, M. Cutadapt removes adapter sequences from high-throughput sequencing reads. EMBnet.journal 17, 10–12 (2011).

69. Bolger, A. M., Lohse, M. & Usadel, B. Trimmomatic: a flexible trimmer for Illumina sequence data. Bioinforma. Oxf. Engl. 30, 2114–2120 (2014).

70. Ramírez, F. et al. deepTools2: a next generation web server for deep-sequencing data analysis. Nucleic Acids Res. 44, W160–165 (2016).

71. Love, M. I., Huber, W. & Anders, S. Moderated estimation of fold change and dispersion for RNA-seq data with DESeq2. Genome Biol. 15, 550 (2014).

72. Alasoo, K. wiggleplotr: Make read coverage plots from BigWig files. (2021).

73. Wang, J., Vasaikar, S., Shi, Z., Greer, M. & Zhang, B. WebGestalt 2017: a more comprehensive, powerful, flexible and interactive gene set enrichment analysis toolkit. Nucleic Acids Res. 45, W130–W137 (2017).

74. Hughes, C. S. et al. Single-pot, solid-phase-enhanced sample preparation for proteomics experiments. Nat. Protoc. 14, 68–85 (2019).

75. Strasser, L. et al. Proteomic Landscape of Adeno-Associated Virus (AAV)-Producing HEK293 Cells. Int. J. Mol. Sci. 22, 11499 (2021).

