## Supplementary material for "Annotation of the non-canonical translatome reveals that CHO cell microproteins are a new class of mAb drug product impurity": Description of Supplementary Data

Supplementary Data 1

**Cell densities at 72 hr post-seeding for the NTS and TS sample groups for both the initiation and elongation experiments.** A significant reduction in cell density was observed in the temperature shifted group when compared to the non-temperature shift group for both experiments.

Supplementary Data 2

**Read pre-processing metrics for Ribo-seq and RNA-seq data.** The number of reads removed following adapter trimming and quality assessment are included for each dataset. For the Ribo-seq data the number of reads eliminated following alignment to contaminating RNA species (rRNA, tRNA and snoRNA) as well as phasing analysis are shown.

Supplementary Data 3

**ORFs identified in this study.** ORF-RATER was used to annotate ORFs using the 3 types of Ribo-seq data. This table contains both annotated and novel ORFs identified. For each ORF, the ORF-RATER ID, ORF-type, ORF-RATER score, gene symbol, gene name, transcript family, transcript ID, start codon, amino acids, whether the start or stop codons were previously annotated, transcript and genome coordinates and strand are included.

Supplementary Data 4

**Host cell protein identification.** LC-MS/MS analysis of host cell protein impurities for **(a)** pertuzumab, **(b)** adalimumab, **(c)** denosumab and **(d)** vedolizumab. Shown are the UniProt and ORF-RATER accession, sequence coverage, number of peptides, length, Sequest score. In addition, the Normalised abundance of biological and technical replicates are shown along the HCP concentrations determined by the Hi3 method are provided for each detected protein.

Supplementary Data 5:

**Significant differences in RNA abundance and translational regulation observed following temperature shift.** DESeq2 was used to identify changes in RNA abundance, RPF occupancy and translational efficiency for **(a-c)** canonical ORFs and **(d-f)** sORFs found in ncRNA. The ID assigned by plastid, the NCBI Gene ID or ORF-RATER ID, gene symbol (for canonical ORFs), Gene Description (for canonical ORFs), baseMean of DESeq2 normalised counts, log_2_ p-value and BH adjusted p-value are shown for each gene/ORF.

Supplementary Data 6

**GO biological process enrichment analysis for differentially translated canonical genes.** The GO ID, description, number of genes in category, overlap, enrichment ratio, p-value, FDR, and genes overlapping are shown for each overrepresented biological process.

Supplementary Data 7

**Temperature shift experiment protein identification.** List of **(a)** NCBI annotated proteins, **(b)** novel ORFs ≥ 100 aa and **(C)** microproteins identified from the temperature shift experiment.

Supplementary Data 8

**Growth phase experiment protein identification.** List of **(a)** NCBI annotated proteins, **(b)** novel ORFs ≥ 100 aa and **(C)** microproteins identified from the growth phase experiment.

Supplementary Data 9

**Temperature shift differentially expressed proteins. (a)** Canonical proteins and **(b)** microproteins identified for the TS24 hr samples vs NTS as well as **(c)** canonical proteins and **(d)** microproteins for the TS48 hr samples vs NTS.

Supplementary Data 10

**Growth phase differentially expressed proteins. (a)** Canonical and microproteins found to be differentially expressed between the Day 4 and Day 7 samples.

Supplementary Data 11

**Database search settings for (a)** cell lysate proteomics and **(b)** drug product HCP analysis**.**
