## Supplementary Results for "Annotation of the non-canonical translatome reveals that CHO cell microproteins are a new class of mAb drug product impurity"

**Upstream open reading frames repress the translation efficiency of CHO cell mRNAs**

The presence of uORFs in 5’ leader sequences has been shown to have a repressive effect on the main ORF in multiple species^1^, and we wished to evaluate if the presence of one or more predicted uORFs or ouORFs, was associated with a change in translation efficiency or RNA abundance. For this analysis, we calculated the average of RPKM values for the CDS of annotated transcripts identified by ORF-RATER (n=7,571), for both the RNA-seq and CHX Ribo-seq data across all replicates of the NTS and TS groups. Transcripts with an average RNA RPKM < 0.5 were eliminated from further analysis, as well as those that contained both an uORF and ouORF. The translational efficiency (TE) for a given transcript, was determined by first dividing the Ribo-seq RPKM by the RNA-seq RPKM for each sample, before the average TE across the 8 replicates was calculated.

The majority of uORF containing transcripts contained a single uORF but, in some cases, as many as 7 uORFs were identified for a single transcript (Supplementary Figure 11a), while 93.5% of transcripts with an ouORF contained a single ouORF (Supplementary Figure 11d). The transcripts were first split into groups based on the number of uORFs or ouORFs present, and the cumulative distribution of both the translation efficiency and RNA abundance for these groups was compared. For uORFs, we stratified the transcripts into those without an upstream ORF (uORF or ouORF) (n=5,882), transcripts containing a single uORF (n =1,974), two uORFs (n=826) or > 3 uORFs (n=388). A significant difference in TE was observed for each uORF containing group when compared to the no-uORF control group, with the degree of repression increasing as the number of uORFs per transcript increased (Figure 11b). We also observed differences in RNA abundance for the uORF containing transcript groups compared to the control group (Supplementary Figure 12a). For ouORFs, we compared ouORF containing transcripts (n=1,125) to the control group. As with uORFs, the ouORF containing transcripts had a significantly reduced translation efficiency (Supplementary Figure 11e), however no impact on RNA abundance was observed (Supplementary Figure 12c). To determine if different uORF or ouORF start codons affect the repression of the main ORF, we focussed on those transcripts that contained a single upstream ORF and compared the cumulative distribution of translation efficiency and RNA RPKM for each of the four start codon groups to the control groups. The AUG start codon in both uORFs and ouORFs was associated with the largest degree of repression compared to the control transcripts, in terms of translation efficiency (Supplementary Figure 11c & Supplementary Figure 11f). We observed a more modest repressive effect on RNA RPKM for transcripts with CUG-, GUG- and UUG-initiating uORFs (Supplementary Figure 11b), while we observed no significant difference between RNA transcripts with near cognate-initiating ouORFs and AUG-initiating ouORFs (Supplementary Figure 11d). There was no significant difference in TE between transcripts containing a single AUG initiated uORF, and those with a single AUG initiated uoORF (Supplementary Figure 13a), although there was a slight difference in RNA abundance between these two groups (Supplementary Figure 13b).

**References**

1.	Chew, G.-L., Pauli, A. & Schier, A. F. Conservation of uORF repressiveness and sequence features in mouse, human and zebrafish. *Nat. Commun.* **7**, 11663 (2016).

### Supplementary Figures

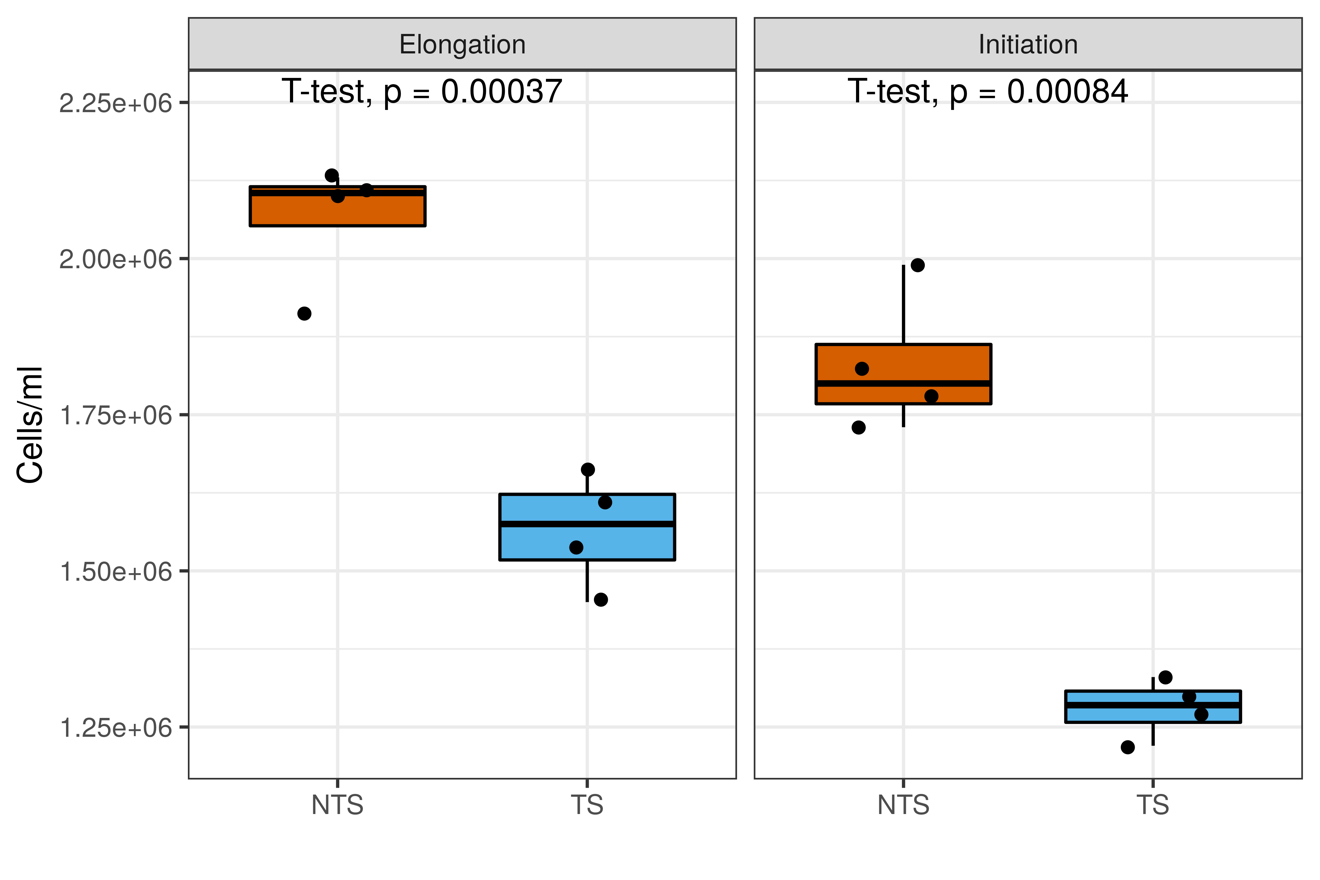

**Supplementary Figure 1: Reduction of cell culture temperature to 31°C decreases the growth rate of the CHO K1-mAb cell line.** Separate cell culture experiments of the temperature shift model were carried out to generate samples for both elongation Ribo-seq (CHX and RNA-seq) and initiation Ribo-seq (HARR and ND). In both experiments, a significant decrease in cell density of ~25% (elongation) and 31% (initiation) for TS samples was observed 24 hr post-temperature shift (72 hr post-seeding).

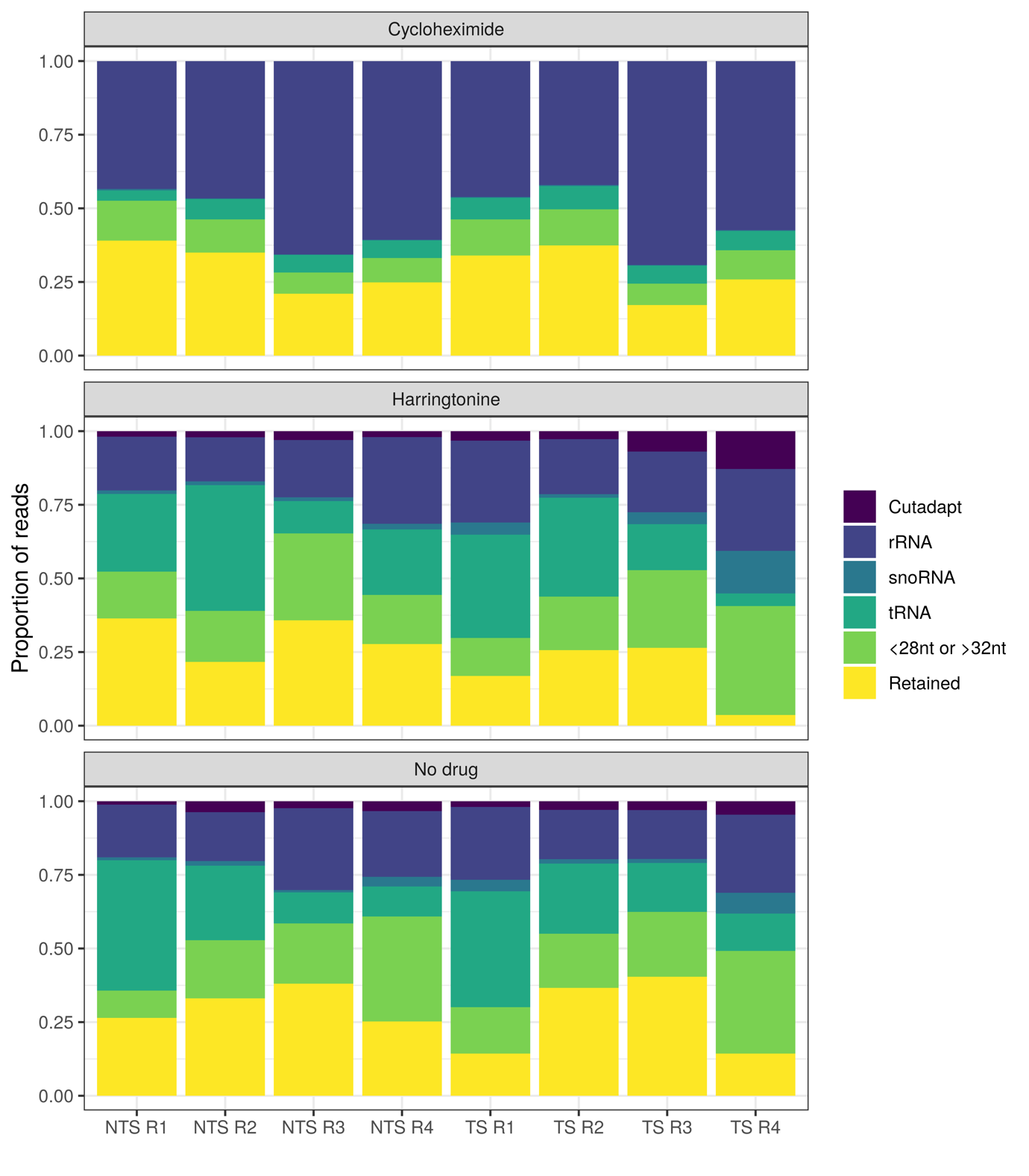

**Supplementary Figure 2: Pre-processing of Ribo-seq data.** Prior to analysis adapters were removed from the raw sequencing reads using Cutadapt. *Note*: The Ribo-seq CHX data was obtained from the sequencing provider with adapter sequences removed. Reads mapping to contaminating RNA species (i.e., rRNA, snoRNA or tRNA) were filtered. Finally, only the reads lengths 28-31nt where 60% of reads were in frame with P-site offset =12, were retained for further analysis.

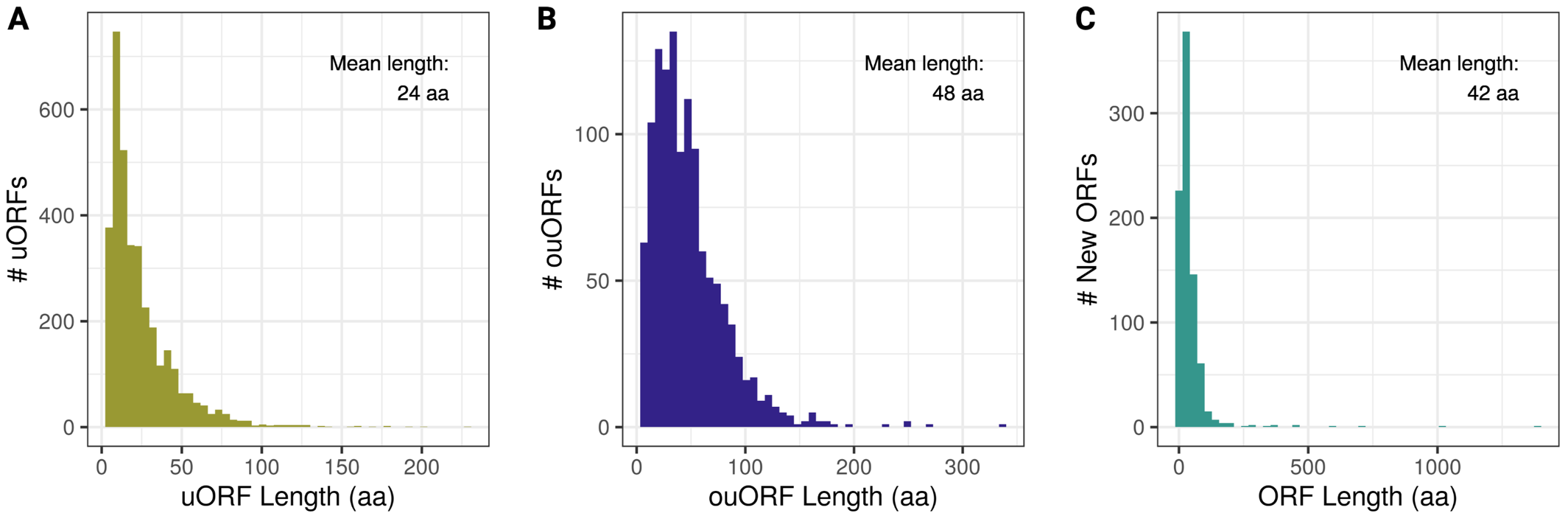

**Supplementary Figure 3: Length distribution of novel Chinese hamster ORF classes comprised primarily of sORFs.** >90% of **(A)** uORFs, **(B)** ouORFs and **(C)** New ORFs identified by ORF-RATER were classified as short ORFs. The average length of each type is shown. Note New ORFs here are found in both protein coding and non-coding transcripts.

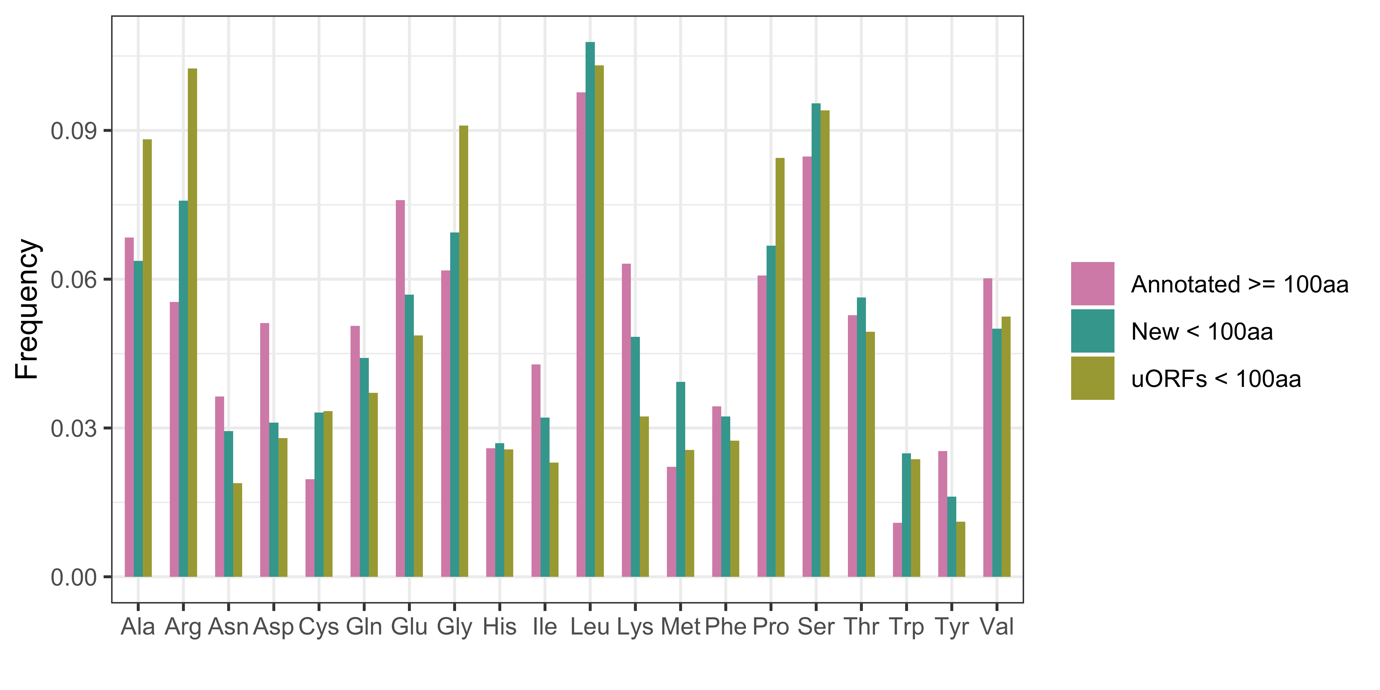

**Supplementary Figure 4**: **Amino acid frequency of annotated and short ORFs.** The amino acid frequencies of the 20aas for uORFs (both uORFs and ouORFs) and ncRNA sORFs along with annotated protein coding genes were determined, revealing differences between each of the groups.

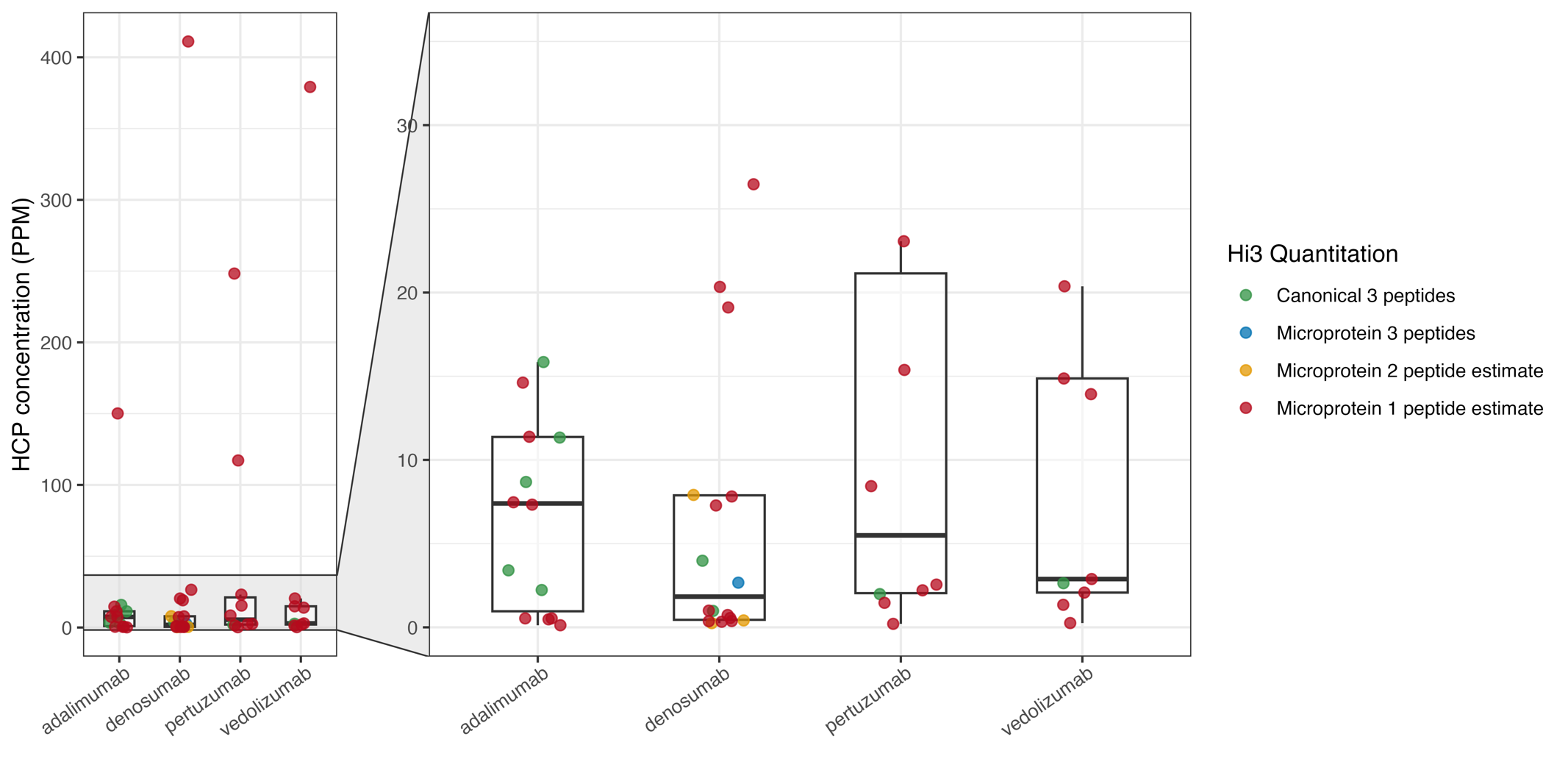

**Supplementary Figure 5: Quantification of microprotein HCPs.** The concentration of canonical proteins and microproteins were determined in 4 commercial antibody drug products using the Hi3 quantitation method. Confident quantitation was achieved for proteins where 3 peptides were identified. The concentrations shown for proteins with 1 or 2 peptide identification should be considered as estimates.

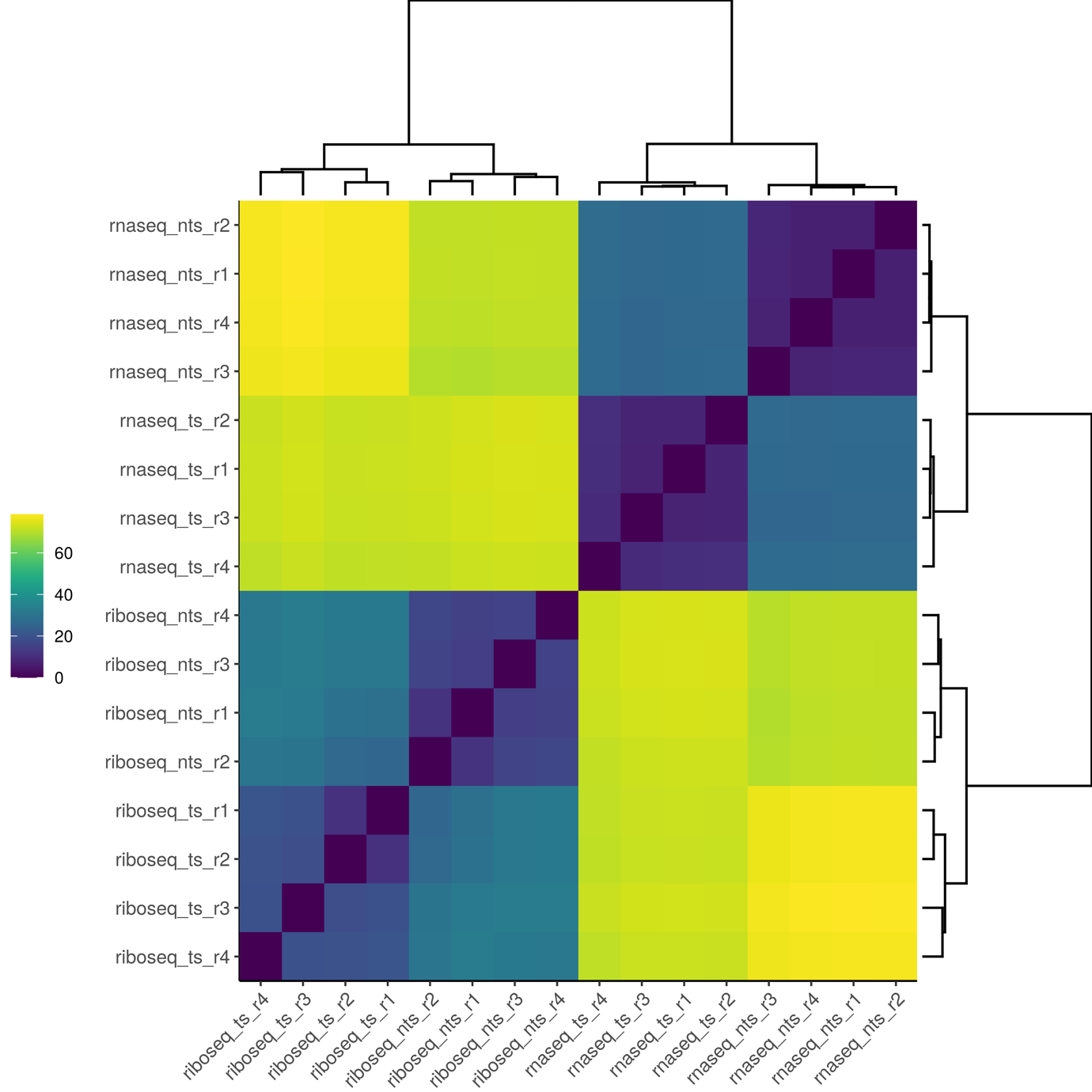

**Supplementary Figure 6: Hierarchical cluster analysis of RNA-seq and CHX Ribo-seq gene-level counts.** While the most significant difference was between the Ribo-seq and RNA-seq data we also observed that there was a clear difference between the NTS and TS sample groups**.**

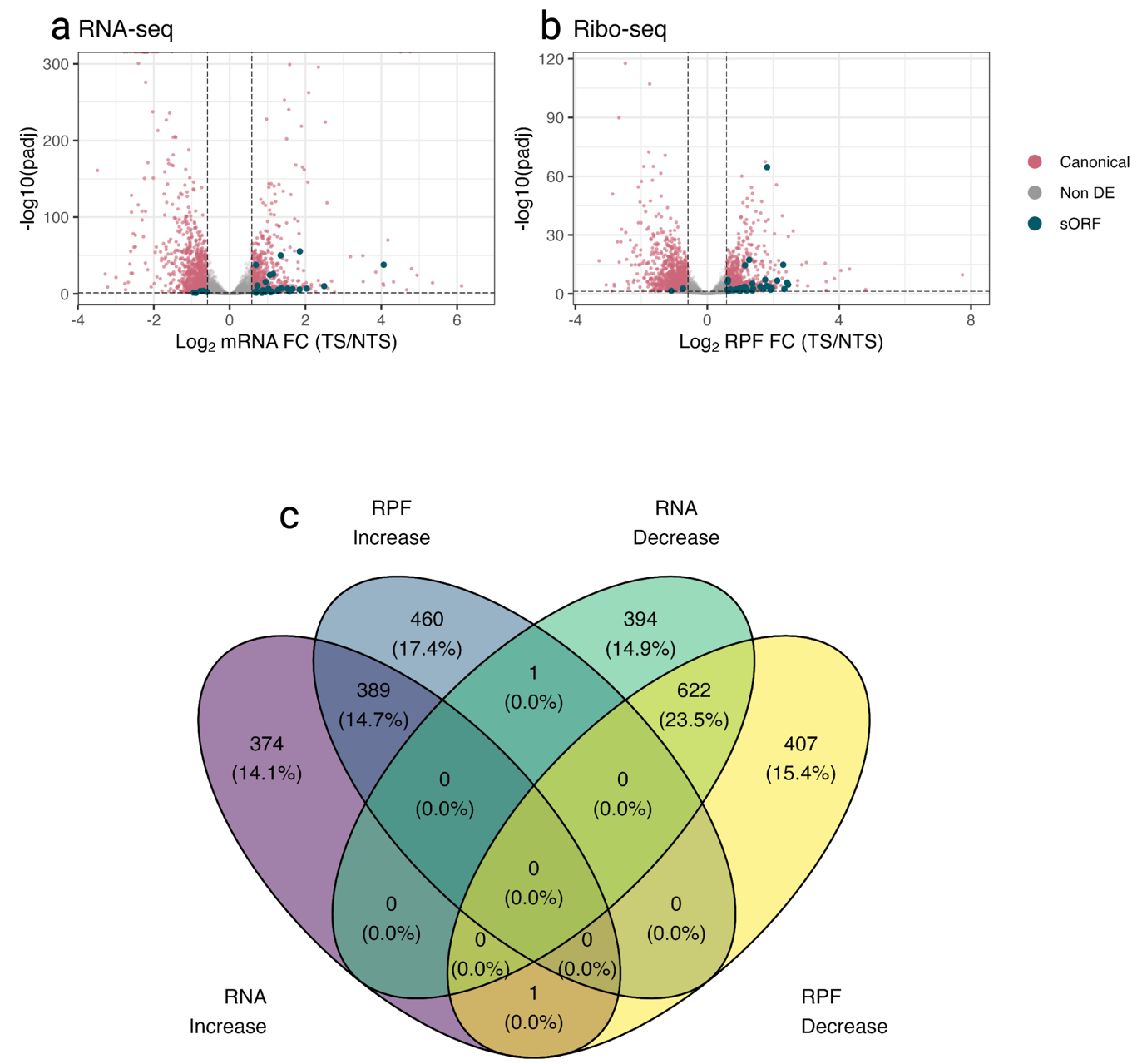

**Supplementary Figure 7: Differential expression and RPF occupancy for canonical and sORFs found in non-coding RNA.** DESeq2 was utilised to identify differences in canonical and sORFs that occurred upon a reduction of cell culture temperature from separate analysis of RNA-seq and Ribo-seq data. A total of **(a)** 1,781 ORFs were found to be differentially expressed from the RNA-seq data and **(b)** 1,880 from the Ribo-seq data. **(c)** 1,011 ORFs were found to change in the same direction in both datasets.

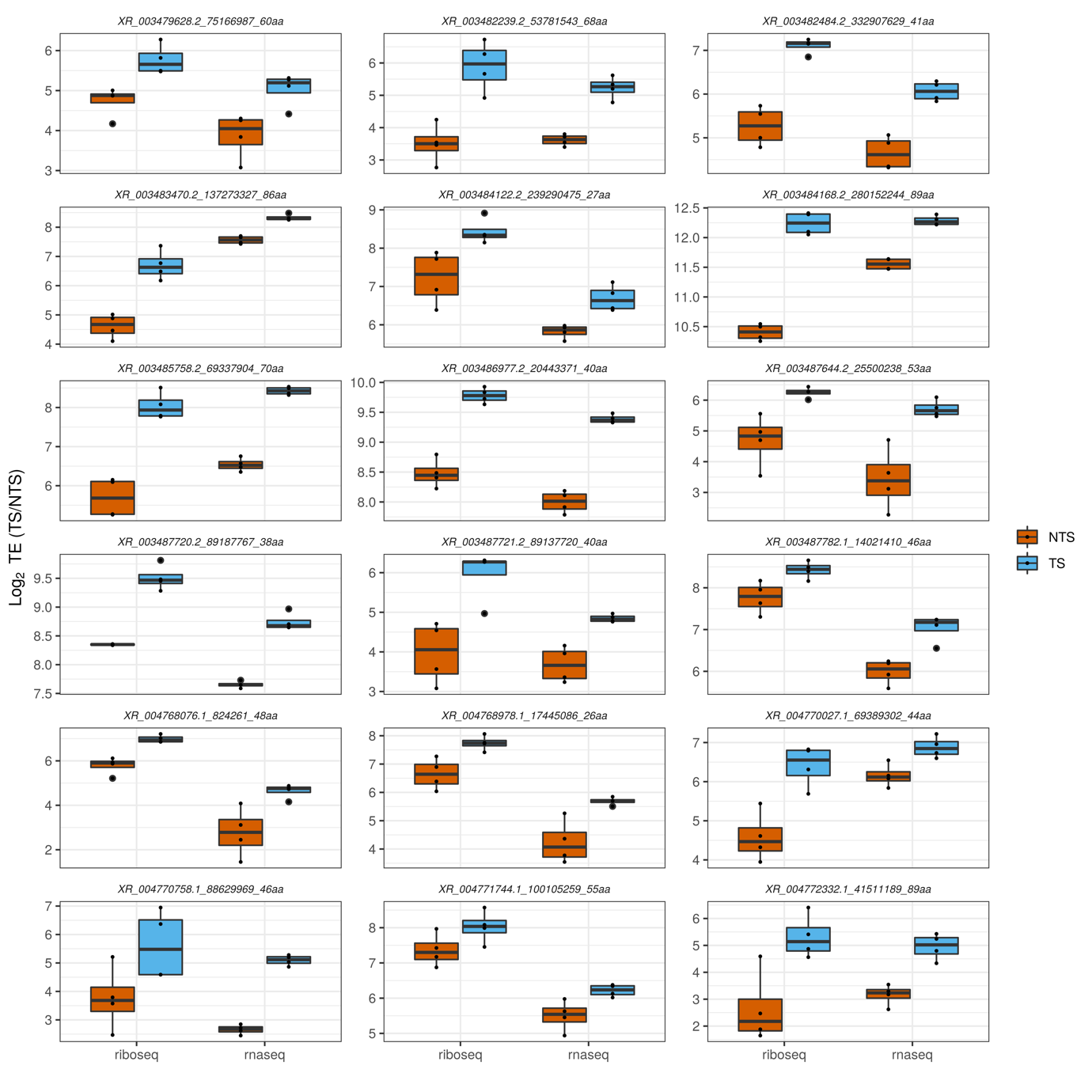

**Supplementary Figure 8: 18 sORFs were found to be upregulated at sub-physiological temperature in both the RNA-seq and Ribo-seq data.**

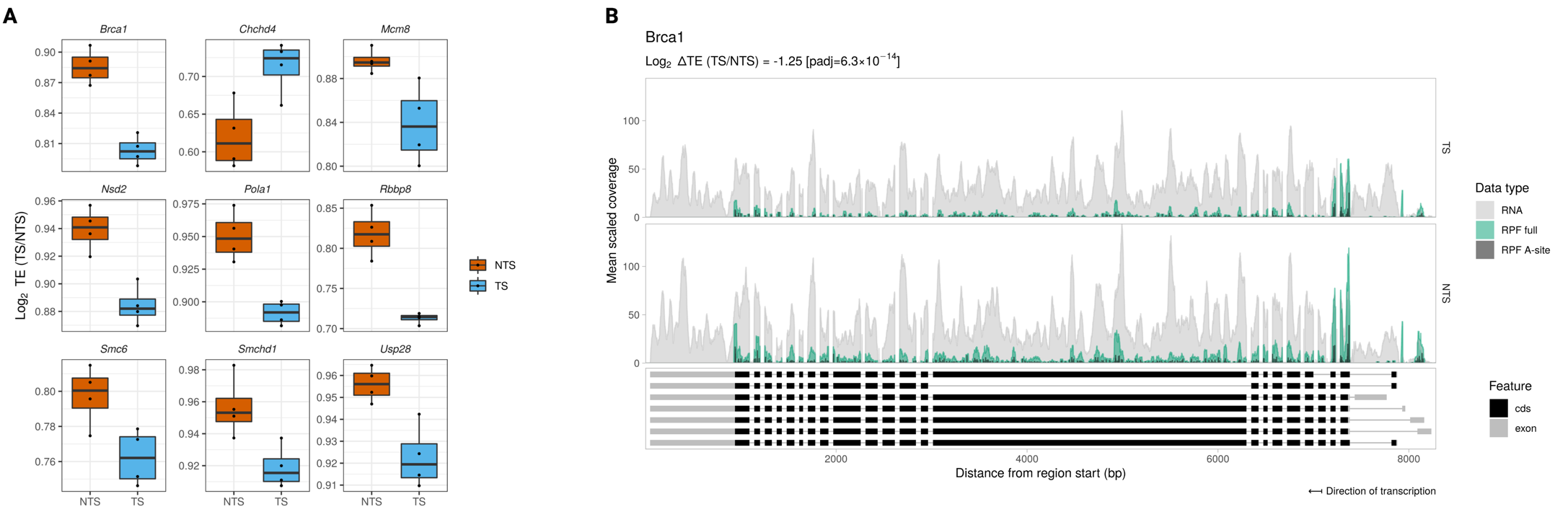

**Supplementary Figure 9: Differential translation efficiency of canonical ORFs involved in DNA repair.** GO enrichment analysis revealed the significant overrepresentation of genes involved in the DNA repair. **(A)** Translation efficiency 9 of the 26 genes related in the DNA repair biological process including **(B)** *Brca1* were found to be altered by a reduction in cell culture temperature.

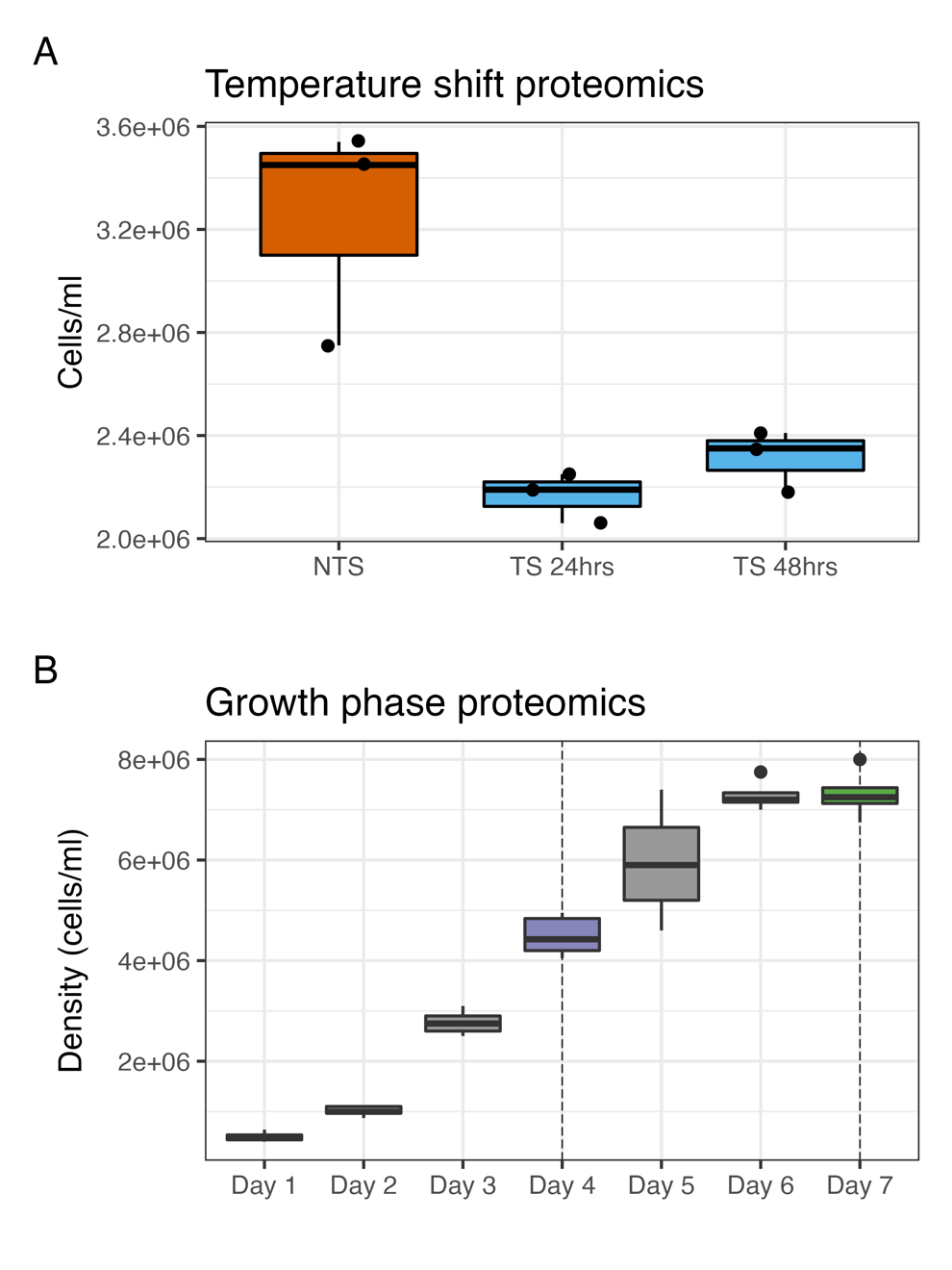

**Supplementary Figure 10: Cell culture conditions used to detect microproteins by mass spectrometry. (A)** We harvested non-temperature shifted cells at 72 hours post seeding. Temperature shifted cells harvested at 24 hours (72 hours post seeding) and 48hours (96 hours post seeding). **(B)** A non mAb-producing CHOK1GS cell line, and captured samples at the exponential growth (Day 4) and stationary phases (Day 7) for proteomics. Proteins were extracted from cell lysates (4 biological replicates for each condition).

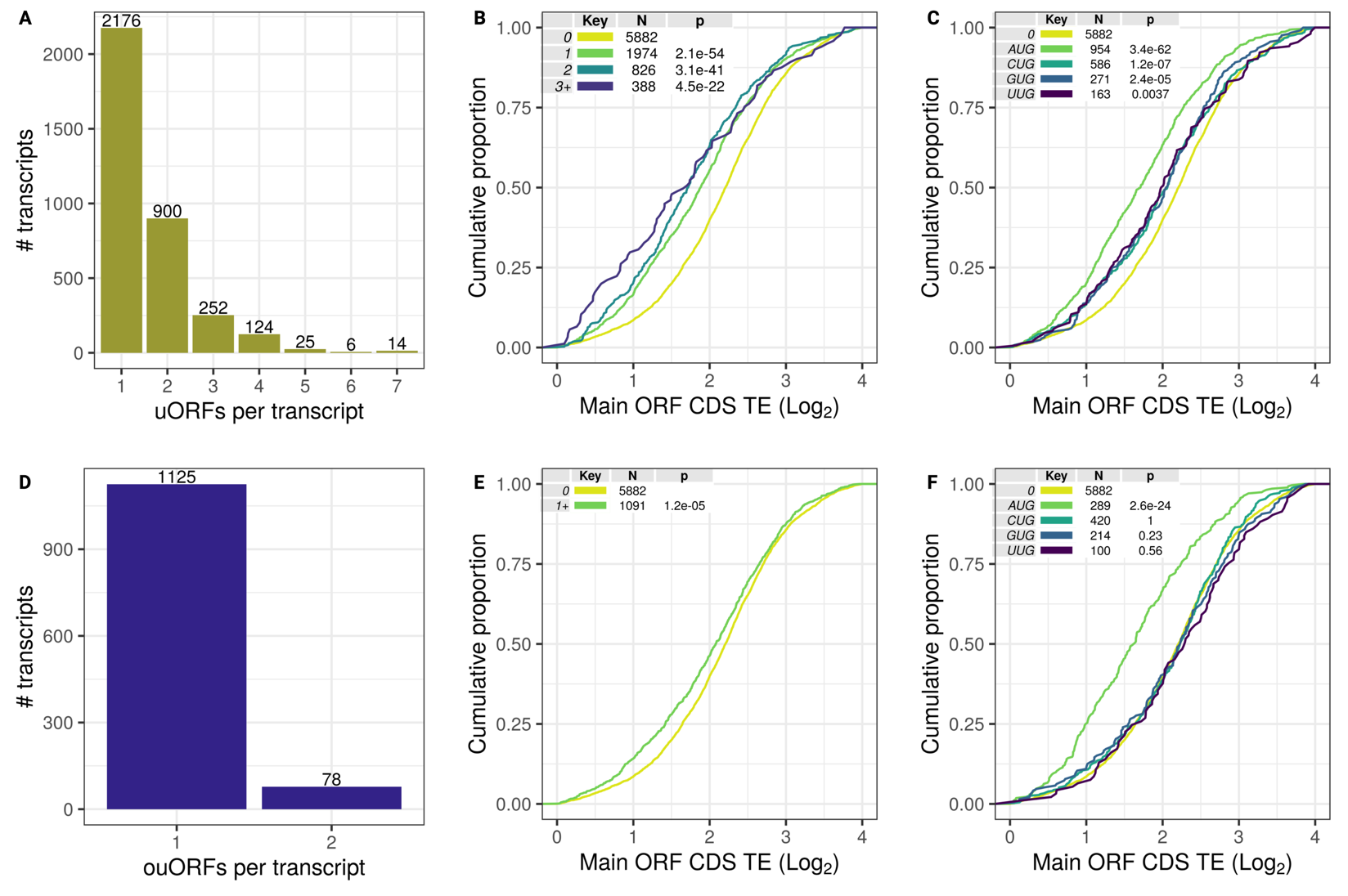

**Supplementary Figure 11: Upstream ORFs tend to repress the translation efficiency of the main ORF.** Ribosome footprint profiling data enabled the identification of translated regions in the 5’ leader sequences of Chinese hamster mRNAs. These upstream ORFs comprised two classes that, were either separate from the main ORF (uORFs) or extended beyond the start codon of the main ORF (ouORFs). To obtain an overview of the transcriptome wide impact of uORFs on the translation efficiency of the main ORF, we stratified annotated protein-coding mRNAs identified by ORF-RATER with an RNA RPKM ≥ 0.5 by the number of uORFs and by start codon for transcripts containing a single uORF**.** For uORFs, most uORF-containing mRNAs had **(A)** 1 or 2 uORFs but for some mRNAs as many as 8 were identified**.** We divided the transcripts into 4 groups (0, 1, 2 and more than 3 uORFs [3+]). There was **(B)** a statistically significant reduction in TE between each of the 3 groups (Wilcoxon test) in comparison to those transcripts without a uORF. To assess the impact of different start codons, only those transcripts with a single uORF were used. When compared with the no-uORF control **(C),** uORFs initiated at each start codon were associated with a significant repression of the main ORF, with the degree of repression of AUG initiated uORFs greater than that of near-cognate start codons. For ouORFs, the **(D**) majority of transcripts contained 1 ouORF, with a small number of instances of 2 ouORFs. We compared all ouORFs containing transcripts [1+] against the control group, and **(E)** a statistically significant impact on TE of the main ORF was observed. Comparison of the start codon for single ouORFs to the control group, revealed **(F)** a significant reduction of TE only for ouORFs starting at AUG.

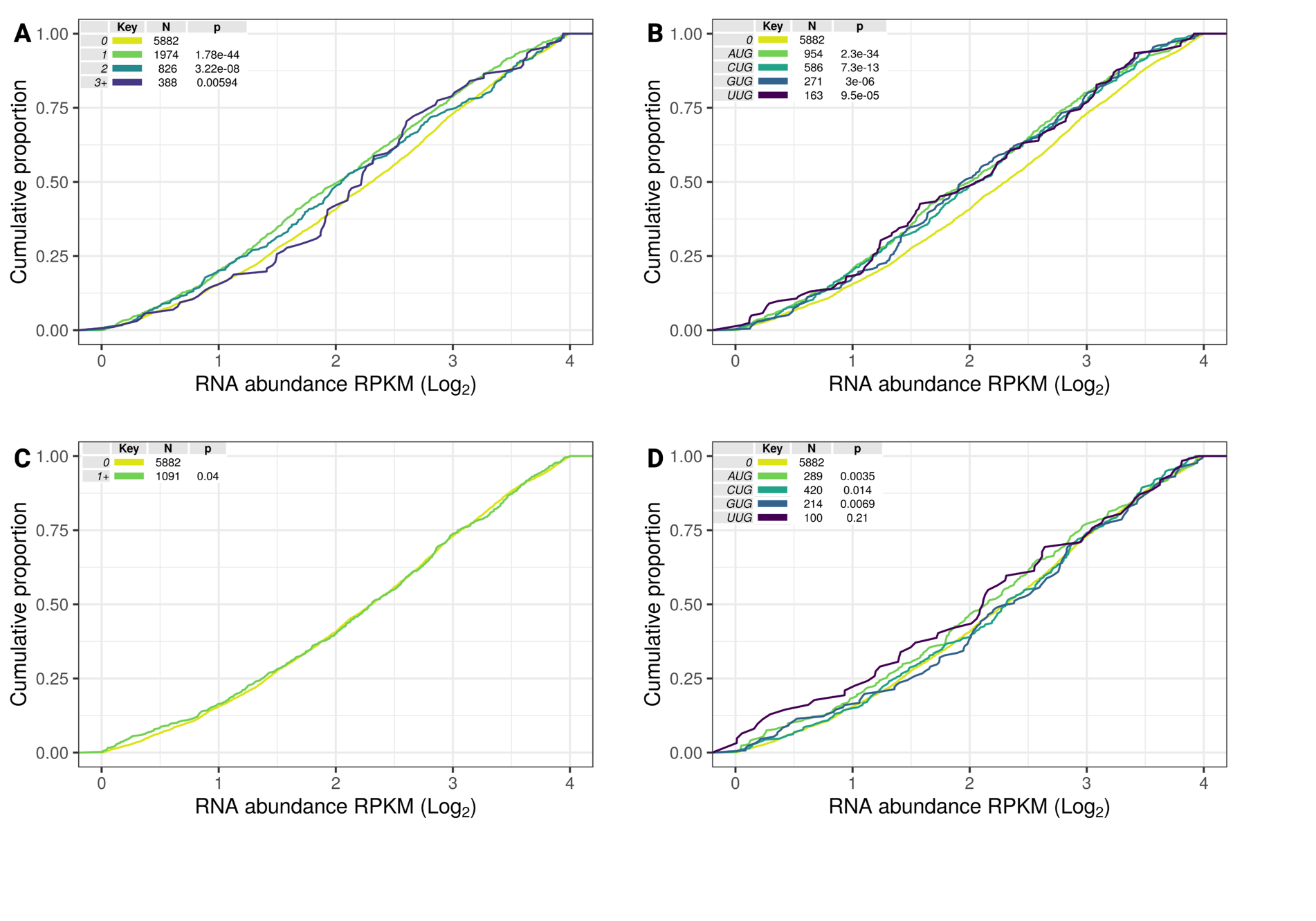

**Supplementary Figure 12: Comparison of uORFs and ouORFs impact on RNA abundance of the main ORF CDS.** For this analysis, in the same manner as the impact of upstream regions on TE transcripts were split into groups based on, either the number of uORF/ouORFs per transcript or transcripts containing a single uORF/ouORFs were dived based on the start codon of the upstream region. This analysis was limited to the annotated transcripts identified by ORF-RATER and the RNA abundance is average RPKM expression from the 8 replicates comprising both the NTS and TS groups. For uORFs, we observed that the **(A)** number that those transcripts containing were both significantly different that the no ouORF control, the 3+ transcript group while statistically significant the difference to the control group was less clear. In the case of single uORF containing transcripts, we observed that **(B)** uORFs initiated at AUG and the near cognate codons were all significantly different in comparison to the no-uORF control. No association between the number of **(C)** ouORFs or **(D)** start codons and RNA abundance was observed.

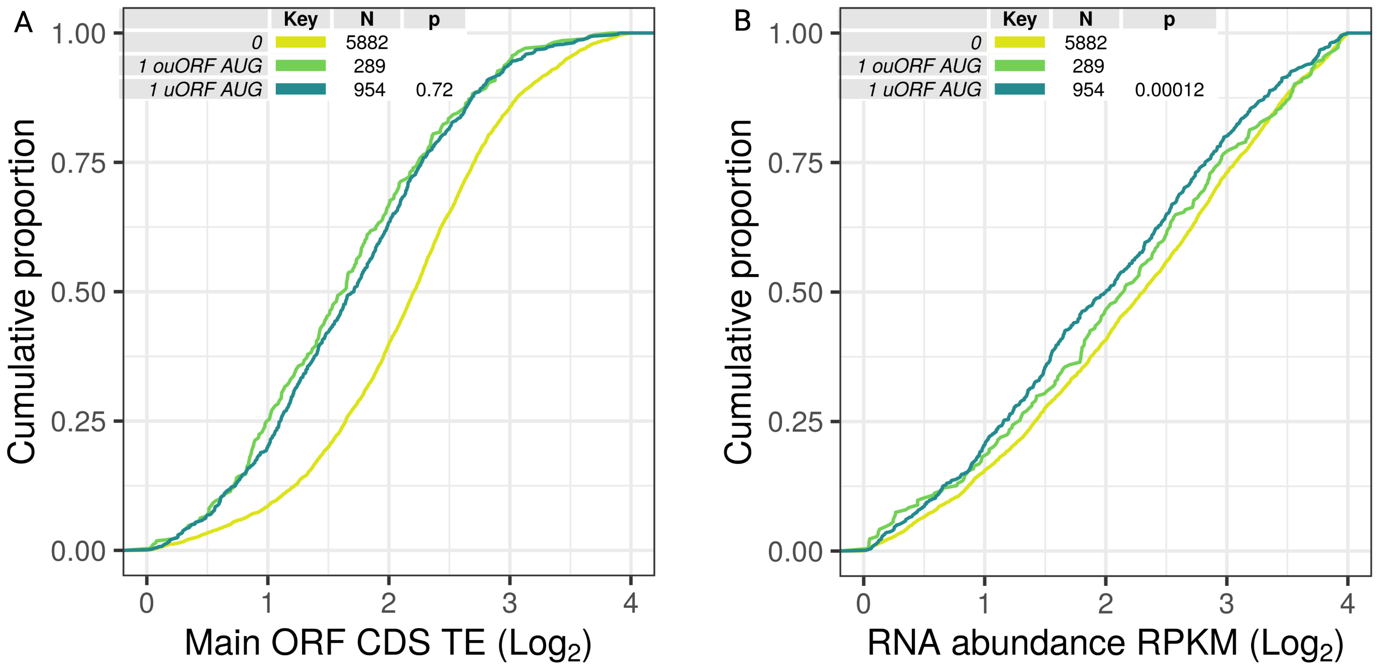

### Supplementary Figure 13: Comparison of AUG initiated uORFs and ouORFs. To compare uORFs and ouORFs, we selected transcripts where a single uORF or ouORFs with an AUG start codon was identified and compared the TE and RNA abundance against a no-uORF/no-ouORF control. The presence of both uORFs and ouORFs significantly reduced (A) the translational efficiency versus the control – however (B) no notable difference in RNA abundance between the uORF and ouORF containing transcripts was observed.
